# Normalised whole-plant water supply declines with plant height across the seed-plant phylogeny

**DOI:** 10.64898/2026.11.18.767533

**Authors:** Victor Flo, Luke Yates, Manon Sabot, Laura Fernández-de-Uña, Jordi Martinez-Vilalta, Nate McDowell, Pablo Sanchez-Martinez, Yanchao Zhao, Huiying Xu, Andrea Carminati, Leander D. L. Anderegg, William R. L. Anderegg, Anna T. Trugman, Maurizio Mencuccini

## Abstract

Whole-plant water-transport capacity largely controls terrestrial water fluxes, yet it is unresolved to what extent its variation across the seed-plant phylogeny is structured by shared ancestry, edaphoclimatic conditions, plant height or water relation traits. We fitted a phylogenetic structural equation model to maximum leaf-specific whole-plant hydraulic conductance ( *k_L_*), together with concurrent Huber value (*Hv*), stomatal conductance (*gs*), and height for 263 species spanning the global climate space, partitioning direct from indirect effects, and phylogenetically conserved from independent effects. *k_L_* declined with height, and *Hv* did not offset this decline. *Hv* raised *k_L_* but was unrelated to height among species. *k_L_* was lower in wetter niches, despite higher *Hv* there, indicating lower conductance per unit sapwood area. *k_L_* was tightly coupled to *gs* in both conserved and independent variation, so the height-driven decline in *k_L_* propagated to gas exchange. Plant height therefore constrains whole-plant hydraulic supply and, through it, *gs*.

## INTRODUCTION

Terrestrial vegetation channels water into the atmosphere at a maximum rate set by the soil-to-leaf hydraulic conductance. Expressed per unit leaf area, whole-plant hydraulic conductance (*k_L_*) defines the capacity of the transport system to supply water for leaf gas exchange, a capacity that is itself strongly modulated by soil water status (Mencuccini 2003). *k_L_* thereby constrains leaf transpiration and photosynthesis and, at larger scales, ecosystem water and carbon fluxes (Binks *et al*. 2022). Species differ in *k_L_* by several orders of magnitude (Mencuccini 2003), yet the factors that structure this variation across seed plants remain unresolved. It is unclear whether species-to-species variation in whole-plant water transport capacity is governed by shared evolutionary history, by the environmental niche species occupy, by plant height (*h*), or by how plants allocate sapwood area relative to leaf area (Huber value, *Hv*; Huber 1928). Because *h* and allocation may themselves be shaped by phylogeny and climate, an important question is whether these factors act independently or in coordination.

Taller plants experience longer hydraulic pathways and, all else being equal, a greater total resistance per leaf area (the inverse of *k_L_*); gravity further lowers the water potential their canopies must sustain. Although path-length resistance is only one component of whole-plant resistance, it underpins the long-standing hypothesis that increasing height lowers *k_L_* and thereby constrains stomatal conductance (*gs*) and carbon assimilation (Mencuccini & Grace 1996; Ryan *et al*. 2006; Ryan & Yoder 1997), with consequences for the drought vulnerability of tall plants (McDowell & Allen 2015). Subsequent work, however, proposed that plants maintain leaf-specific supply as they grow taller by widening their conduits, increasing *Hv*, or both (compensation hypothesis). Conduits widen predictably from stem tip to base, and basal conduits are wider in taller plants across angiosperms (Olson *et al*. 2014, 2018). Under hydraulic optimality models, this tapering is expected to compensate for path length by holding *k_L_* approximately constant as plants grow taller (Lechthaler *et al*. 2020; Petit & Anfodillo 2009). A higher *Hv* could likewise hold leaf-specific water supply relatively constant (Becker *et al*. 2000). Because wider conduits raise sapwood-specific conductivity (*K_s_*), taller species also tend to have higher *K_s_*, particularly in wet climates, leading to the proposal that height and hydraulic traits form a single coordinated axis aligned with water availability (Liu *et al*. 2019).

Most of this evidence comes from scales below the whole plant, for example from bordered pits that link individual xylem conduits, which alone account for over half of xylem hydraulic resistance (e.g. Choat *et al*. 2008), single xylem conduits (e.g. Olson *et al*. 2014), or organs such as leaves, roots or branches (e.g. Martínez-Vilalta *et al*. 2002). Although these data have contributed to a general understanding of how hydraulic architecture responds to environmental gradients, inferences drawn at one scale are not necessarily valid at another, because selection acts on the integrated organism and it is the combination of whole-plant traits that confers fitness (Meinzer *et al*. 2010).

Where whole-plant conductance has actually been measured, tapering does not fully compensate for height, but this evidence comes largely from ontogenetic series within species, in which conductance declines as individuals grow taller (Martínez-Vilalta *et al*. 2007; Mencuccini & Magnani 2000; Phillips *et al*. 2002; Zaehle 2005). Among species, where the processes involved may differ, the relationship has been examined only in small compilations across life forms (Mencuccini 2003). The same gap applies to the coupling between supply and gas exchange. At the leaf scale, stomata can track hydraulic supply to keep leaf water potential within safe limits (Brodribb & Jordan 2008; Sack & Holbrook 2006), but whether whole-plant *k_L_* and *gs* remain coordinated across species, so that a height-driven decline in supply would reach gas exchange, has rarely been tested beyond that same compilation (Mencuccini 2003).

Given that maximum *h* and *Hv* covary with climate (Mencuccini *et al*. 2019; Scheffer *et al*. 2018), and given that organ-level hydraulic traits may exhibit a substantial phylogenetic signal (Sanchez-Martinez *et al*. 2020), the effects of *h*, *Hv*, climate and shared ancestry on *k_L_* could be confounded. Whether the compensation hypothesis holds for *k_L_* among species, once these effects are jointly resolved, remains untested.

In this study, we examined how *h*, *Hv*, climate and shared ancestry relate to one another and jointly shape *k_L_* across >250 plant species spanning the global climate space (Fig. 1), using observations of maximum *k_L_* from a global database (Zhao *et al*. 2026). We combined *k_L_* with *h*, *Hv*, concurrent stomatal conductance (*gs*), and species-level edaphoclimatic descriptors in a phylogenetic structural equation model (Thorson & Van Der Bijl 2023). Our model partitions the direct from the indirect components, and the phylogenetically conserved from the phylogenetically independent components of the relationships linking *h*, *Hv*, climate, and water transport. We address three questions. First, does *k_L_* decline with *h* through a direct effect that persists after accounting for *Hv* and climate? Because tip-to-base conduit widening appears to be near-universal (Olson *et al*. 2014), any negative effect of height on *k_L_,* once *Hv* is accounted for, would indicate that widening and allocation together do not offset the increase in path length. Based on previous within-species evidence, we expected a negative, allocation-independent effect of height on *k_L_*. Second, does a height-driven decline in *k_L_* propagate to *gs*? Given the coordination between water supply and stomatal regulation, we expected *gs* to decline with height through *k_L_* (Meinzer 2002; Mencuccini 2003). Third, are the relationships among these traits phylogenetically conserved or expressed independently of shared ancestry? Because *k_L_* characterises a maximum capacity, and *gs* the conductance concurrent with it, both are expected to evolve slowly under environmental filtering and stabilising selection rather than track short-term meteorological variability. We therefore expected their coordination to be phylogenetically structured (Sanchez-Martinez *et al*. 2024).

**Figure 1.**
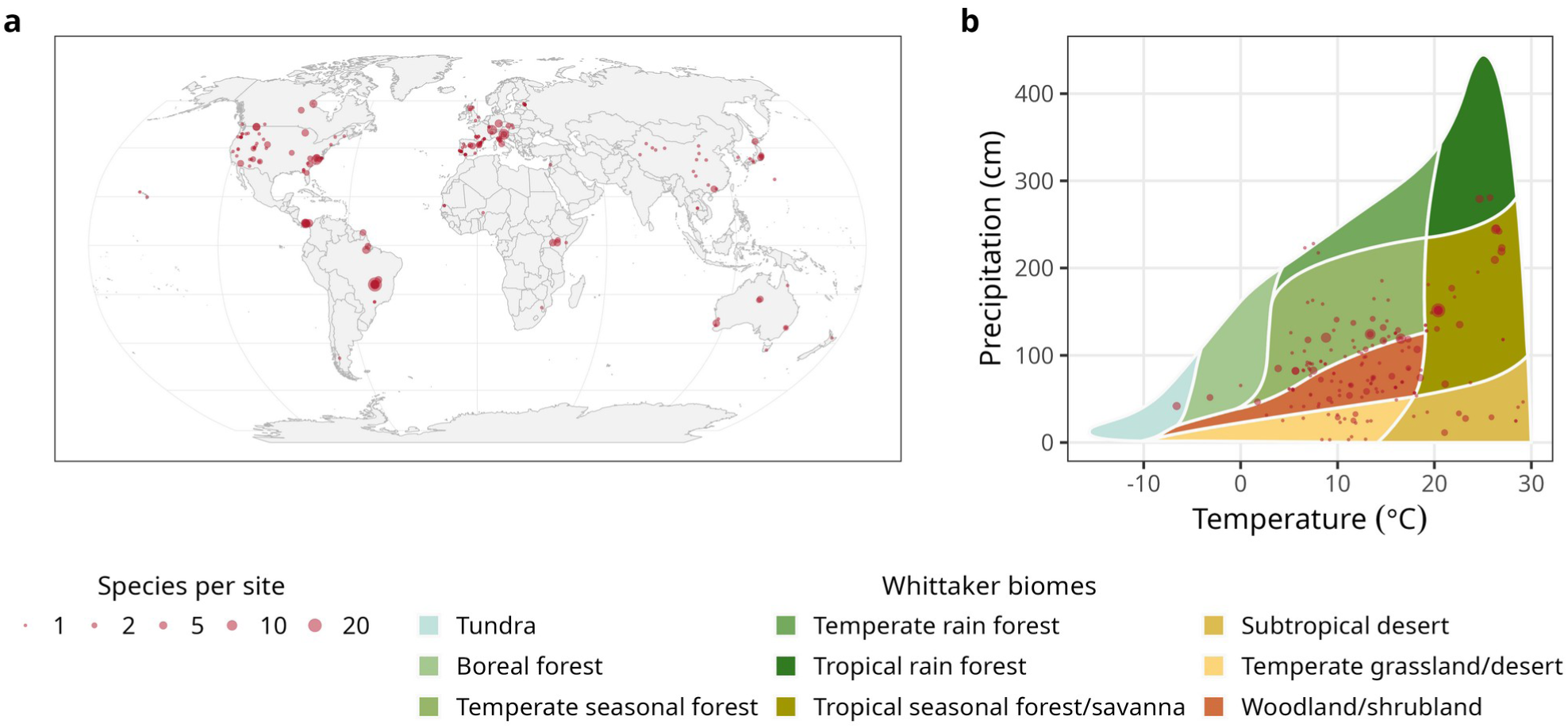
Geographic and bioclimatic distribution of the data. (a) Global distribution of the 194 sites included in the analysis. (b) Distribution of sites in Whittaker’s climate space (mean annual temperature vs. mean annual precipitation; biome boundaries after Whittaker 1975). Dot size indicates the number of species recorded at each site.

## MATERIALS AND METHODS

### Data sources

We extracted four plant variables, leaf-specific whole-plant hydraulic conductance (*k_L_*, kg m^-2^ s^-1^ MPa^-1^), plant height (*h*; m), Huber value (*Hv*) and stomatal conductance (*gs*, mol m^-2^ s^-1^), from the KPLANT database (Zhao *et al*. 2026), restricting the selection to field-grown adult plants not subjected to experimental manipulation (Table S1). For each species at each site, we selected the record with the highest *k_L_* across all individuals and measurement dates, and took *h*, *Hv* and *gs* from that same record, so that the four variables describe a single individual. Species-site values were then averaged across sites to obtain one estimate per species. The final dataset comprised 263 species from 194 sites (Fig. 1). We treat the resulting *k_L_* as operational estimate of its species-level maximum and *gs* as the conductance concurrent with it rather than as a species-level maximum. *Hv* was mostly estimated at the whole-tree level (86% of source studies), predominantly from sapwood area at breast height (69%). Because *h* refers to the measured individual rather than to the species, we assessed its representativeness against species-level maximum heights from TRY (95th percentile of non-experimental records; Kattge *et al*. 2020).

For each species, we characterised the climatic and edaphic niche using occurrence records from GBIF (Global Biodiversity Information Facility (GBIF) 2026). Occurrences were cleaned using the CoordinateCleaner package (Zizka *et al*. 2019), removing duplicates, coordinates matching country capitals or biodiversity institutions, records falling in the sea, and zero or equal coordinates. Geographic outliers within each species were then removed with a quantile-based test, flagging records whose geographic distance to all other records of the species exceeded the upper quartile by more than five interquartile ranges. Two species (*Saccharum hybr* and *Eucalyptus sclerophylla*) with fewer than seven unique localities were not tested for outliers. Similarly to Sanchez-Martinez *et al*. (2020), at each retained occurrence we extracted ten climatic variables from WorldClim v2.1 at 30 arc-second resolution (Fick & Hijmans 2017): six standard bioclimatic variables (mean annual temperature, annual precipitation, temperature seasonality and precipitation of the warmest quarter, wettest month and driest quarter), together with four variables derived from monthly layers: mean annual maximum temperature, mean annual solar radiation, maximum monthly wind speed, and the mean vapour pressure deficit of the warmest month, calculated as the difference between the mean saturation vapour pressure, averaged over the monthly minimum and maximum temperatures, and the actual vapour pressure. We additionally used an aridity index from the CGIAR-CSI database (Zomer *et al*. 2022) and five variables from SoilGrids 2.0 at 1 km resolution (Poggio *et al*. 2021): pH and sand, silt and clay fractions, each averaged over the 0-200 cm profile with weights proportional to layer thickness, and volumetric water content at field capacity (-33 kPa) at 100-200 cm (Turek *et al*. 2023), as a measure of deep-soil water retention. Sand, silt and clay fractions were expressed as two isometric log-ratio coordinates (sand relative to silt and clay, and silt relative to clay; Muzzamal *et al*. 2018) to account for their compositional nature. We computed the species-level mean across occurrences for each edaphoclimatic variable and summarised the fifteen variables using a principal component analysis (PCA). We retained the first three components (PC1-PC3), which together explained 83.8% of the total variance (Fig. S1). PC1 defined an axis of water availability and temperature, with positive values associated with high annual precipitation and, to a lesser extent, high mean annual temperature, and negative values with high temperature seasonality. PC2 represented an axis of atmospheric demand, with positive values associated with higher vapour pressure deficit of the warmest month and radiation. PC3 reflected a soil-texture gradient, with positive values associated with sandier soils and negative values with higher silt and clay content. Species PC scores were merged with the trait dataset.

### Phylogeny

We reconstructed the phylogenetic relationships among the 263 species in the dataset using V.PhyloMaker2 (Jin & Qian 2022), using the backbone GBOTB.extended mega-phylogeny of seed plants (Smith & Brown 2018). 201 of our species matched taxa already present in the tips of the backbone phylogeny, while the remaining were inserted at the genus level (56 species) or at the family level following scenario S3 (6 species; Jin & Qian 2022). The resulting tree was made ultrametric using the “extend” method within phytools 2.0 (Revell 2024).

### Statistical analysis

Prior to species aggregation and analysis, *k_L_*, *h*, *Hv* and *gs* were log-transformed to (i) make their multiplicative inter-relationships additive in the linear model, and (ii) to stabilise the residual variance (Fig. S2). All variables entering the phylogenetic structural equation model were additionally standardised to achieve zero mean and unit variance, thus path coefficients represent standardised effect sizes (Gonzalez-Voyer & Von Hardenberg 2014). Trait coverage was uneven across the 263 species (Fig. S3): PC1-PC3 and *k_L_* were available for all the species, whereas *h*, *gs* and *Hv* were only available for 147, 126, and 90 species, respectively. Missing trait values were treated as model parameters and imputed jointly with the model parameters, conditional on the phylogeny and the path structure.

We modelled the network linking species-average edaphoclimatic conditions, plant size, allocation, hydraulic transport and stomatal conductance as a phylogenetic structural equation model. Each species’ K-vector of traits (or edaphoclimatic variables) was modelled as:

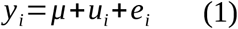

where *i*=1 , … *, N* indexes species, *μ* is the vector of ancestral means, *ui* the phylogenetically conserved component and *ei* the phylogenetically independent component, including measurement error.

The conserved components are correlated among species according to phylogeny such that cov (*ui , u j* )=*V ij Σ*phy where *V* is the correlation matrix implied by Brownian motion on the phylogeny rescaled to unit root-to-tip height. The phylogenetically independent components are independent among species cov (*ei , e j* )=*δij Σ*ind, where *δᵢⱼ* equals 1 if i = j and 0 otherwise. Each component has its own trait covariance matrix (*Σ_phy_* for the conserved and *Σ_ind_* for the phylogenetically independent component), and it is on these matrices that the causal structure is imposed.

Within each component *c* (conserved or phylogenetically independent), the traits follow a recursive path model in causal order, so that covariation among traits takes the structural form (Bollen 1987; Thorson & Van Der Bijl 2023):

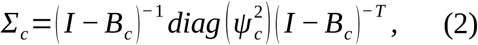

where *B_c_* is the matrix of path coefficients, which is strictly lower-triangular and non-zero only on the retained directed acyclic graph edges (dropped edges are fixed to zero). The *ψ ^2^* are the disturbance variances of individual equations, that is the variance in each trait not accounted for by its parents in the network.

Two path structures were fitted. A full model (m7) specified PC1-PC3 as predictors of *k_L_*, *h*, *Hv* and *gs*; *H* as predictor of *Hv*; *H* and *Hv* as predictors of *k_L_*; and *k_L_* as a predictor of *gs* (Fig. 2, Table 1). A reduced model (m5) retained only PC1-PC3 and *h* as predictors of *k_L_*. This reduced structure isolates the *h*-*k_L_*relationship from sparser downstream traits and, since *h* is the only sparse trait it contains, can be fitted to complete cases as an imputation-free counterpart to m7.

**Figure 2.**
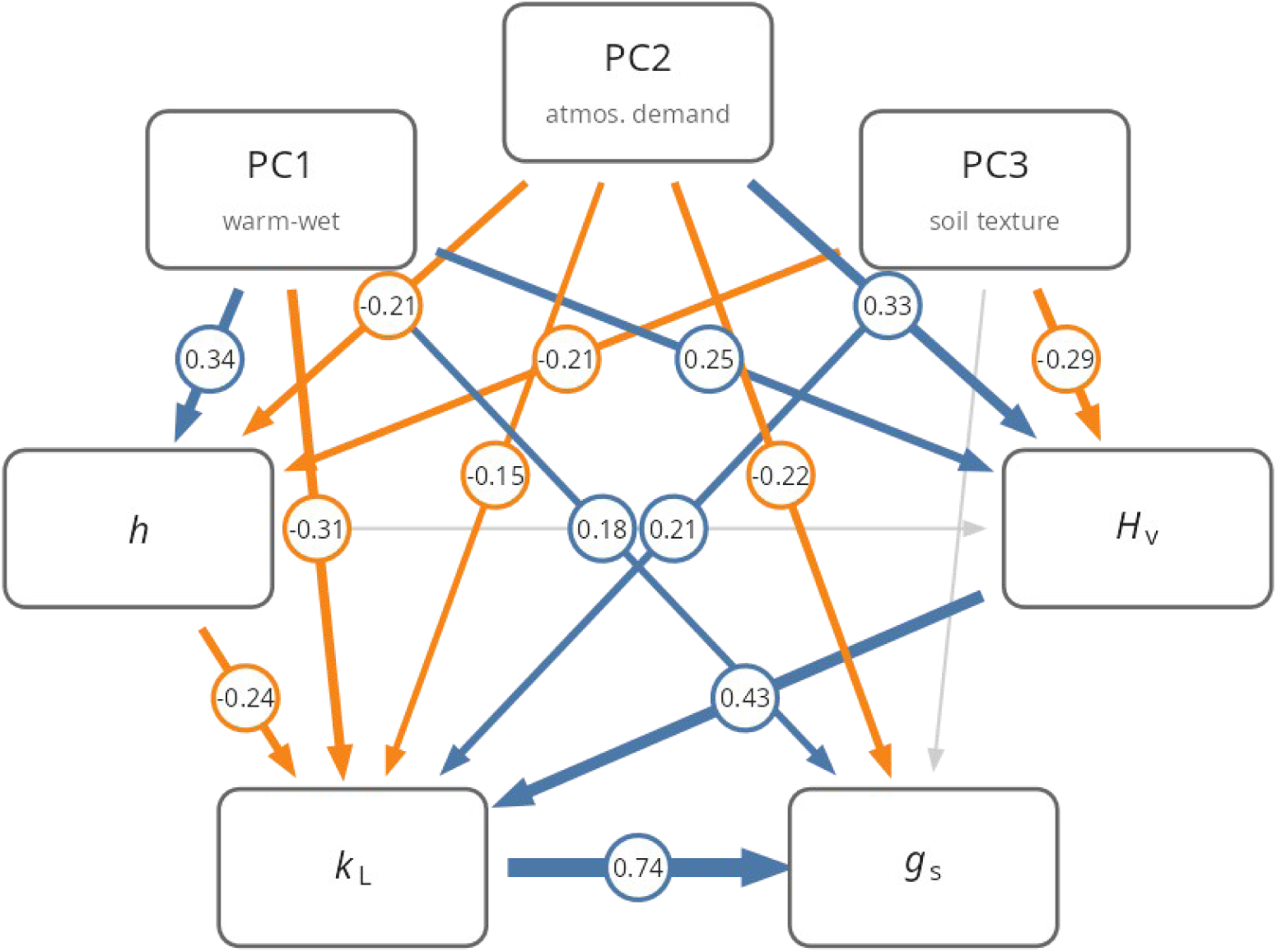
Network of the s*hared form* m7 model showing the direct and indirect paths linking edaphoclimatic axes (PC1-PC3), *h* and *Hv* to *k_L_* and *gs*. Blue arrows denote paths whose 95% credible interval excludes zero and is positive, orange denotes negative paths and grey paths whose interval overlaps zero; arrow width scales with the absolute standardised network coefficient shown alongside each path. PC1 increases toward warm, wet, low seasonality niches; PC2 toward high VPD and solar radiation; PC3 toward sandier soils.

**Table 1.** Hypothesised paths in the m7 network, with rationale and supporting literature.

| Path | Rationale | References |
|---|---|---|
| PC1-PC3 → *h* | Plant height is pedo-climatically constrained. Water availability and thermal regime set an upper limit on canopy height, with soil texture modulating water supply. | (Liu *et al*. 2019; Tao *et al*. 2016) |
| PC1-PC3 → *Hv* | Sapwood-to-leaf allocation acclimates and adapts to water availability and evaporative demand; drier climates favour higher *Hv*. | (Mencuccini & Grace 1996; Sanchez-Martinez *et al*. 2020) |
| PC1-PC3 → *k<sub>L</sub>* | There is direct climatic selection on *k<sub>L</sub>* beyond that mediated by size and allocation. Organ-level evidence predicts a positive effect, with more conductive xylem favoured by warm, wet environments, and hydraulic safety features emerging in dry climates. Because *k<sub>L</sub>* integrates root, stem and leaf resistances whose relative contributions may vary with pedo-climate, we do not assume that organ-level responses transfer linearly to the whole-plant. | (Liu *et al*. 2019; Sanchez-Martinez *et al*. 2020) |
| PC1-PC3 → *gs* | Climate may affect *gs* beyond its coordination with hydraulic supply, for instance through evaporative demand and water availability acting on stomatal regulation. Alternatively, maximum stomatal conductance may converge across climates, in which case climatic effects on *gs* would be transmitted entirely through *k<sub>L</sub>*. | (Lin *et al*. 2015; Murray *et al*. 2019) |
| *h* → *Hv* | Compensation predicts higher *Hv* in taller plants. Alternatively, stretching predicts lower *Hv*, as conducting area per unit leaf area decreases with path length. | (Becker *et al*. 2000; Couvreur *et al*. 2018) |
| *h* → *k<sub>L</sub>* | Hydraulic-limitation prediction: longer transport paths and gravitational potential reduce leaf-specific whole-plant conductance | (Mencuccini 2003) |
| *Hv* → *k<sub>L</sub>* | For a given xylem permeability and path length, a higher Huber value raises the water supply available per unit leaf area. | (Mencuccini *et al*. 2019) |
| *k<sub>L</sub>* → *gs* | Stomata regulate aperture to keep xylem tension above cavitation thresholds, so stomatal conductance measured concurrently with maximum *k<sub>L</sub>* tracks the hydraulic supply available per unit leaf area. Neither *h* nor *Hv* has a direct mechanistic link to *gs*; both act on stomatal behaviour through the hydraulic supply they generate. We orient the path from supply to regulation because *k<sub>L</sub>* changes over longer time scales than *gs*, but species-level trait covariation cannot establish direction, and demand-driven investment in transport capacity would produce a comparable association. | (Carminati *et al*. 2026; Meinzer 2002) |

Each path structure was fitted in two forms. In the s*hared form,* a single path network enters both components, so that the conserved, phylogenetically independent and phenotypic-total networks coincide and only the disturbance variances differ between components (Hardenberg & Gonzalez-Voyer 2013; Thorson & Van Der Bijl 2023; Van Der Bijl 2018). In the s*eparated form*, *B_phy_* and *B_ind_* were estimated independently, allowing path coefficients to differ in magnitude and sign between the conserved and phylogenetically independent components (Notes S1) (Shipley 2009). The s*eparated form* is a constrained submodel of a multi-response phylogenetic mixed model (Hadfield & Nakagawa 2010; Halliwell *et al*. 2025).

The phylogenetic signal per trait was summarised in two complementary ways: a per-equation fraction *λeq*, the share of a trait’s disturbance variance attributable to the phylogenetically conserved component, and a marginal fraction *λmarg*, the conserved share of its total modelled variance (equivalent to Pagel’s λ (Pagel 1999), see also Note S1). The two definitions coincide for the exogenous PCs and diverge for downstream traits, where conserved variation additionally reaches a trait through the fitted paths (Table 2).

**Table 2.** Phylogenetic signal per trait from the s*hared form* m7 model on the full dataset (N = 263), given in two ways: *λ_eq_*, the fraction of the variance of a trait not explained by its predictors in the network (its residual variance) that is phylogenetically conserved, and *λ_marg_*, the fraction of its total modelled variance that is conserved once signal propagated through the network is included (Note S1). Values are posterior means with 95% credible intervals. PC1-PC3 are the edaphoclimatic axes; *Hv*, Huber value; *k_L_*, whole-plant leaf-specific hydraulic conductance; *gs*, stomatal conductance.

| Trait | *λ<sub>eq</sub>* | *λ<sub>marg</sub>* |
|---|---|---|
| PC1 | 0.93 [0.88, 0.97] | 0.93 [0.88, 0.97] |
| PC2 | 0.82 [0.69, 0.90] | 0.82 [0.69, 0.90] |
| PC3 | 0.24 [0.03, 0.53] | 0.24 [0.03, 0.53] |
| *h* | 0.71 [0.47, 0.87] | 0.75 [0.59, 0.87] |
| *Hv* | 0.29 [0.00, 0.73] | 0.46 [0.18, 0.74] |
| *k<sub>L</sub>* | 0.66 [0.31, 0.86] | 0.67 [0.50, 0.80] |
| *gs* | 0.19 [0.01, 0.48] | 0.46 [0.29, 0.62] |

To separate the direct and mediated contributions of each predictor we decomposed the fitted network into direct, indirect, and total effects on *k_L_*. The direct effect is the path coefficient linking one trait to another, i.e. the part of their association not routed through any intermediate trait in the network. The total effect additionally accumulates every directed route linking the two traits. It is obtained from the coefficient matrix as (I - B)^-1^ - I (see Note S1.6). We computed each indirect effect as the difference between the total and direct effects. All three quantities were computed on every posterior draw, so uncertainty propagates through the products and sums of coefficients, and the effects are reported as posterior means with 95% credible intervals given by the 2.5th and 97.5th percentiles. In the s*eparated form*, the decomposition was applied to *B_phy_*and *B_ind_* independently, yielding conserved and phylogenetically independent direct, indirect and total effects for every trait pair.

Whole-plant conductance can be estimated from different whole-plant flux data, one of which is sap flow (Zhao *et al*. 2026). Because the proportion of sap-flow-based *k_L_* estimates covaried with plant height, potentially confounding the *h*-*k_L_*relationship with the measurement method, we refitted the model separately to species whose maximum *k_L_* included sap-flow-based estimates (N = 135). Trait availability also depended on the method used to estimate *k_L_*, of the 90 species with an observed *Hv*, 80 came from records including sap flow, so no independent check on the *Hv* paths was possible. The estimates also differed in whether *k_L_* and *gs* were instrumentally independent. Where *k_L_* derives from sap flow, *gs* is measured separately (N = 72), whereas methods based on evaporative fluxes typically derive both *k_L_* and *gs* of the same transpiration measurement (N = 54). The sap-flow subset therefore provides the estimate of the *k_L_*-*gs* coupling least susceptible to shared-measurement inflation. To test for potential effects related to plant growth form or taxonomic groups, we further refitted the model to trees alone (N = 192) and to angiosperms alone (N = 232), gymnosperms (N = 31) were too few to be analysed separately. We refitted the reduced model (m5) to species with complete observations for all its variables (N = 147), providing an imputation-free counterpart. Because *h* was measured on the sampled plants, it reflects their size at the time of measurement rather than species potential stature. To test whether results depended on this, we refitted the m7 network replacing measured *h* with species maximum height from the TRY database (Kattge et al. 2020; TraitIDs 18 and 3106). For each species, we took the 95th percentile of height records, excluding records flagged as minimum or low values and records from experimental conditions (207 species). Measured *h* and TRY maximum height were correlated (r = 0.73, n = 123), with sampled plants reaching a median of 37% (range 10-76%) of maximum height. Standardisation and phylogenetic tree rescaling were recomputed within each subset.

All models were fitted by Hamiltonian Monte Carlo in Stan (Carpenter *et al*. 2017), using CmdStan 2.38 through cmdstanr (Gabry *et al*. 2025), with data preparation and post-processing done in R 4.3.3 (R Core Team 2024). We rotated the species onto eigenvectors to factorise the joint likelihood into N independent K-variate normal densities for computationally efficient sampling (Note S1.2).

Priors were weakly informative: N(0, 1) on path coefficients, half-Cauchy(0, 5) on disturbance standard deviations, and N(0, 10) on ancestral means (Note S1). Each model was run as four chains of 4000 iterations, including 2000 iterations discarded as warm-up, with adapt_delta = 0.95 and a maximum tree depth of 12. Convergence was assessed from split-R^ (all <1.01) and bulk effective sample sizes (all > 400) (Vehtari *et al*. 2021). We assessed model adequacy with two posterior predictive checks of species-level *k_L_*, each shown against the observed values. The first predicts a species from its parents in the fitted network, the second from the remainder of the dataset (Note S1.7, Fig. S4).

## RESULTS

The fitted m7 s*hared form* network using all the available data was densely connected (Fig. 2), and the edaphoclimatic axes were related to almost all traits. Species occupying warm, wet niches with low seasonality (high PC1) were taller (0.34 [0.19, 0.49]) and had higher *Hv* (0.25 [0.03, 0.49]) but lower *k_L_* (-0.31 [-0.46, -0.15]). Species from niches with high atmospheric demand and radiation (high PC2) were shorter (-0.21 [-0.36, -0.05]), with higher *Hv* (0.33 [0.12, 0.54]) and lower *g_s_* (-0.22 [-0.39, -0.06]). Species in sandier soils with lower deep-soil water retention (high PC3) were shorter (-0.21 [-0.34, -0.07]), with lower *Hv* (-0.29 [-0.53, -0.05]) but higher *k_L_* (0.21 [0.07, 0.35]).

Beyond these edaphoclimatic effects, *k_L_* declined with *h* (-0.24 [-0.43, -0.05]; Fig. 2, Fig. S5), so that taller species had lower *k_L_*. This association was stronger in the reduced model fitted to complete cases, without imputation (m5; -0.31 [-0.50, -0.11]; Fig. S6). The effect of *h* on *k_L_* was not mediated by allocation, as *h* and *Hv* were unrelated (-0.11 [-0.41, 0.19]), leaving no indirect route through *Hv* (Fig. 3). *Hv* did, however, exert an expected positive direct effect on *k_L_* (0.43 [0.23, 0.57]; Fig. S5), since all else being equal, more conductive tissue for a given leaf area increases the water flux supplied per unit of transpiring surface (see Note S2). Finally, *k_L_* was strongly and positively coupled to *gs* (0.74 [0.54, 0.94]; Fig. S5), a coupling credible in both the conserved and the phylogenetically independent components of the s*eparated form* fit (0.73 [0.22, 1.21] and 0.71 [0.39, 1.03], respectively; Fig. S7).

**Figure 3.**
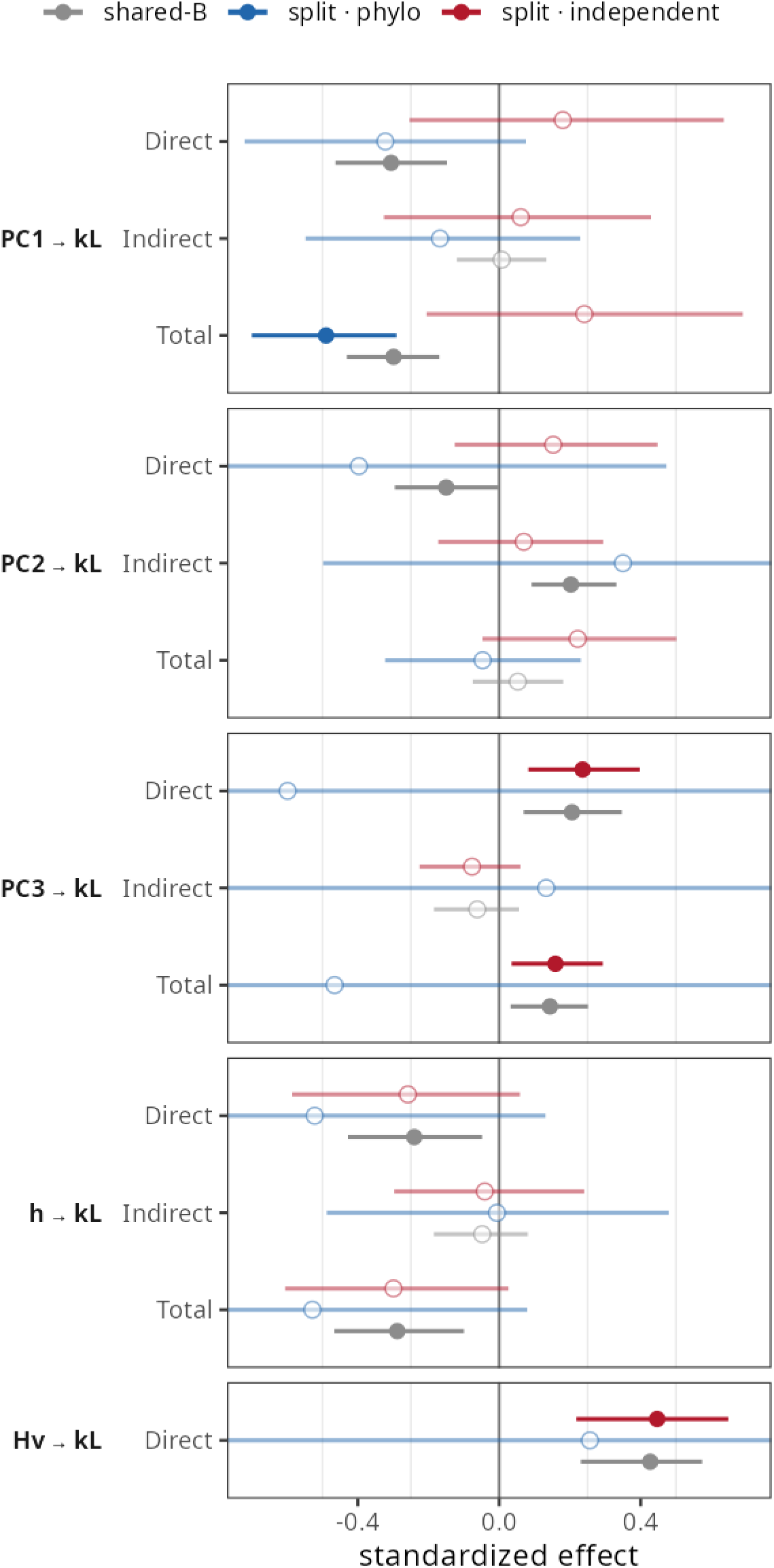
Direct, indirect and total standardised effects of *h*, *Hv* and the edaphoclimatic axes on *k_L_*, decomposed per predictor. Each estimate is shown for three variants of the full model: the s*hared form* network (grey), and the phylogenetically conserved (blue) and phylogenetically independent (red) networks of the s*eparated form* fit. Points show the posterior mean and bars the 95% credible interval; filled points denote effects whose interval excludes zero, open points those overlapping zero. Indirect and total effects are shown only for predictors with a mediated route to *k_L_*. The x-axis is clipped at ±0.7 for readability. PC1 increases toward warm, wet, weakly seasonal niches; PC2 toward high atmospheric demand and radiation; PC3 toward sandier soils.

The trait paths were largely robust to measurement method and to species composition. In species whose maximum *k_L_* included sap-flow-based estimates (N = 135; Fig. S8), *k_L_* still declined with *h* (-0.22 [-0.42, -0.02]) and *Hv* retained its positive effect on *k_L_* (0.52 [0.29, 0.70]). Note, however, that this subset contains 80 of the 90 species with an observed Hv, so it does not provide an independent test of this path. The *k_L_*-*gs* coupling remained strong (0.52 [0.22, 0.81]) and because *gs* is measured independently of *k_L_* wherever *k_L_* derives from sap flow, we regard this as the more conservative estimate of the *k_L_*-*gs* coupling. In trees (N = 192; Fig. S9), the total effect of h on *k_L_* was preserved (-0.20 [-0.37, -0.02]) but its direct component was not credible (-0.13 [-0.30, 0.04]), while the effect of *Hv* (0.56 [0.39, 0.71]) and the *k_L_*-gs coupling (0.67 [0.44, 0.89]) were preserved. In angiosperms (N = 232; Fig. S10), the effect of *h* (total -0.28 [-0.48, -0.07]; direct -0.22 [-0.43, 0.00]), the effect of *Hv* (0.39 [0.12, 0.57]) and the *k_L_*-gs coupling (0.75 [0.52, 0.97]) were preserved. h and *Hv* remained unrelated in all subsets (Table S5). When measured *h* was replaced by TRY maximum height, the total effect of *h* on *k_L_* was preserved (−0.29 [−0.43, −0.14]), although its direct component was not credible (−0.18 [−0.36, 0.01]; Table S5).

To separate the routes underlying the associations revealed by the phylogenetic structural equation models, we decomposed each total effect into direct and mediated paths (Fig. 3). In the s*hared form*, the effect of *h* on *k_L_*was direct rather than mediated by allocation. The indirect path via *Hv* was negligible (-0.05 [-0.19, 0.08]), and the total effect (-0.29 [-0.47, -0.10]) was similar to the direct one. In the s*eparated form* the effect was of similar or larger magnitude, but its credible interval overlapped zero in both the conserved (-0.52 [-1.18, 0.13]) and the independent components (-0.26 [-0.59, 0.06]; Fig. 3, S7). In the s*hared form*, this decline propagated to *gs* through *k_L_* (-0.21 [-0.37, - 0.07]).

Edaphoclimatic effects on *k_L_* were mostly direct (s*hared form*; Fig. 3). PC1 lowered *k_L_*directly (-0.31 [-0.46, -0.15]; Fig. 3), an effect driven by precipitation rather than temperature. In an auxiliary regression of log *k_L_* on the underlying climatic variables, annual precipitation was the only significant predictor (-0.37, p < 0.001; Table S3). PC1 also acted on *k_L_*through two indirect routes of opposite sign (Table S4). Species from high-PC1 niches were taller, which lowered their *k_L_* (PC1 → h → *k_L_*: -0.08 [-0.17, -0.01]), but also had higher *Hv*, which raised it (PC1 → *Hv* → *k_L_*: 0.11 [0.01, 0.22]). Because these routes cancelled out, the total effect of PC1 on *k_L_*(-0.30 [-0.43, -0.17]; Fig. 3) was almost identical to its direct effect. In the *separated form*, this total effect was credible in the conserved component (-0.49 [-0.70, -0.29]) but not in the phylogenetically independent one (0.24 [-0.21, 0.69]; Fig. S7). The climatic effects on *h* were likewise conserved (PC1: 0.39 [0.16, 0.62]; PC2: -0.43 [-0.76, -0.13]), with no credible counterpart in the independent component (Fig. S7). PC2 lowered *k_L_* directly in the s*hared form* (-0.15 [-0.30, -0.00]; Fig. 3), but this was offset by indirect routes through higher *Hv* (0.14 [0.04, 0.26]) and lower *h* (0.05 [0.004, 0.11]; Table S4), so the total effect of PC2 was not credible (0.05 [-0.08, 0.18]; Fig. 3). PC3 increased *k_L_* both directly (0.21 [0.07, 0.35]) and in total (0.14 [0.03, 0.25]; Fig. 3).

The partition of each trait’s residual variance into conserved and phylogenetically independent components in the *shared form* allowed us to quantify how much of each trait’s variation was phylogenetically structured (Table 2). The edaphoclimatic axes PC1 and PC2 carried strong phylogenetic signal, as expected when closely related species occupy similar climatic niches (Losos 2008), whereas the soil axis PC3 was weakly conserved. Plant *h* was strongly conserved, and remained so at the equation level (*λ_eq_* = 0.71), indicating phylogenetic structure in stature beyond the shared edaphoclimatic niche. *k_L_* showed intermediate signal, almost entirely generated in its own equation (*λ_eq_* = 0.66; *λ_marg_* = 0.67). In contrast, the signal of *Hv* and particularly of *gs* was largely propagated through the network. *gs* was weakly conserved in its own equation (*λ_eq_* = 0.19) but reached *λ_marg_* = 0.46 through its coupling to *k_L_* and the edaphoclimatic axes, a coupling of of near-identical strength in the conserved and phylogenetically independent components (Fig. S7). Phylogenetic signal was lower when the analysis was restricted to trees or to angiosperms (*λ_marg_* of *k_L_* = 0.43 and 0.48, respectively; Table S2), consistent with the exclusion of the deepest phylogenetic contrasts in these subsets.

## DISCUSSION

We found that *k_L_* under non-limiting water conditions declines with *h*, supporting the hydraulic-limitation hypothesis (Ryan & Yoder 1997) across species and at the global scale. *Hv* did not offset the decline in *k_L_* with *h*. Although *k_L_* rose with *Hv*, this rise reflects the arithmetic of leaf-specific supply rather than a compensation effect (see below). Moreover, *Hv* was unrelated to *h* among species, so taller species did not build proportionally more sapwood per unit leaf area to compensate for their longer transport path. In contrast, Mencuccini et al. (2019) reported a weak negative relationship between branch *Hv* and maximum *h* across a larger set of species, although maximum *h* was compiled from independent databases rather than measured on the sampled plants, as here. Within species, however, *Hv* increased with *h* (Note S2), as commonly reported during tree growth (McDowell *et al*. 2002).

We found that *k_L_* was tightly coupled to concurrent *gs*, the strongest link in the network, and remained among the strongest where the two traits were measured independently (0.52 [0.22, 0.81]). This coupling is sub-isometric, meaning plants have proportionally lower *gs* per unit *k_L_* at high *k_L_* (Note S2). The coordination between hydraulic conductance and *gs* is well documented at the leaf scale (Brodribb & Jordan 2008; Sack & Holbrook 2006). At the whole-plant scale, however, this coupling has rarely been examined across species (but see Mencuccini 2003; Sabot *et al*. 2020). Our results establish this coupling in a large, phylogenetically explicit sample, where it held at near-identical strength in both the conserved and the phylogenetically independent components (Fig. S7).

The negative *h*-*k_L_* relationship can be understood in the context of the physics of water transport: hydraulic resistance accumulates with path length, so in the absence of any compensatory adjustment, leaf-specific supply would decline in proportion to height, with an exponent of -1 (Mencuccini 2003; Note S2.1), while complete compensation would give an exponent of 0. Because *k_L_* was computed from water-potential differences measured at the same height (predawn vs. midday) or from height-corrected soil water potential, the gravitational component is already removed, and this expectation concerns frictional resistance only. We found a slope much shallower than -1 (-0.42 in angiosperms, -0.44 across all species; -0.27 for the same relationship estimated net of climate and allocation in the m7 network; Note S2). Notably, these slopes agree with the -0.47 obtained by Mencuccini (2003) across life forms, from a compilation an order of magnitude smaller. Part of the compensation likely lies in xylem anatomy: conduits widen from the leaves towards the stem base, which slows the accumulation of resistance along the stem (Lechthaler *et al*. 2020; Olson *et al*. 2014; Zaehle 2005). Because *k_L_*integrates roots, stem and leaves, part of the compensation may also lie in root or leaf xylem (Binks et al. 2022).

*Hv* and *k_L_* were measured independently, but by definition *k_L_* = *k_S_*·*Hv*, where *k_S_*is whole-plant hydraulic conductance per unit sapwood area, so that log *k_L_*= log *k_S_* + log *Hv*. A log-log slope of *k_L_*on *Hv* would be one if *k_S_* were independent of *Hv*. Across the 90 species with an observed *Hv*, the log-log slope was 0.25 (SE 0.10, p = 0.017), equivalent to -0.75 (SE 0.10) for *k_S_* on *Hv* (Notes S2.3), which implies that species building more sapwood per unit leaf area transport proportionally less water through each unit of sapwood. This whole-plant pattern is consistent with the negative relationship between branch xylem-specific conductivity (Ks, conductance per unit cross-sectional area and multiplied by segment length) and branch *Hv* reported across species (Mencuccini et al. 2019; Sanchez-Martinez et al. 2020).

Environmental conditions structured the network partially through plant *h* (Fig. 2). The edaphoclimatic axes were moderate predictors of *h*, with taller species in warm and wet climates with low seasonality and shorter species under high atmospheric demand and sandy soils, consistent with previous reporting on the global controls of maximum tree height (Liu *et al*. 2019; Tao *et al*. 2016, Scheffer *et al*. 2018). The environmental effect on *k_L_*, by contrast, was largely direct and ran counter to the organ-level expectation outlined in Table 1. Among species of similar *h* and *Hv*, those from wetter niches had lower *k_L_*, and this effect of PC1 reflected precipitation rather than temperature (Table S3). Because wetter-niche species also had higher *Hv*, they appear to build more sapwood per unit leaf area, but sapwood that transports less water per unit of its own cross-section. This inverts the expectation carried over from branch-segment measurements, where hydraulic efficiency generally increases with water availability (Liu *et al*. 2019; Sanchez-Martinez *et al*. 2020), and suggests that efficiency measured on a stem segment is not necessarily a good proxy of efficiency over the whole pathway (Binks *et al*. 2022; Meinzer *et al*. 2010). Because *Hv* was itself measured on the stem, this per-area efficiency refers to the stem cross-section rather than to the whole conducting system. The direct effect could be (i) a genuine whole-plant property, with wet-niche species allocating proportionally more resistance to extra-stem tissues, or (ii) a compositional signature, since dry and seasonal niches are occupied disproportionately by ring-porous and drought-deciduous species with higher maximum *k_L_*. The direct effect of PC1 on *k_L_* was credible in all m7 fits (Table S5). It was weaker in the reduced model, which omits *Hv* and therefore absorbs the positive route through *Hv* into the direct path, and in species with complete observations the climatic signal on *k_L_* ran instead through *h* (Table S5). For PC2, the route through *Hv* offset the direct effect, leaving no net effect of atmospheric demand on *k_L_* (Table S4).

Allocation also showed an unexpected pattern. Species in warmer, wetter climates supported more sapwood per unit leaf area (Fig. 2), an effect that reflected temperature rather than precipitation (Note S2.3). This goes against the expectation that higher temperatures, by lowering water viscosity and raising hydraulic conductivity, should reduce the sapwood area needed to supply a given leaf area (Xu *et al*. 2026). It also goes against the global decline of *Hv* with increasing mean annual temperature, precipitation and moisture index (Mencuccini *et al*. 2019). In that global analysis, however, the direct climatic effect on *Hv* vanished once co-varying leaf and xylem traits were included, with the climatic signal running instead through those traits. It is plausible that our model, which does not resolve specific leaf area (SLA) or *Ks*, may retain as direct an effect that a more complete trait network would route indirectly.

The phylogenetic signal we recover for *k_L_* (*λ_marg_*= 0.67, Table 2) is higher than that reported for branch-level leaf-specific conductivity across the seed-plant phylogeny (*λ* = 0.43; Sanchez-Martinez et al. 2020). Integrating the whole root-to-leaf pathway does not dilute the evolutionary conservatism of hydraulic supply as measured at the branch level. Such intermediate signal is typical among hydraulic traits (Liu *et al*. 2015; Maherali *et al*. 2004; Sanchez-Martinez *et al*. 2020) and, in our predominantly woody sample, consistent with a history in which lineages diversify adaptively yet remain bounded by phylogenetic niche conservatism, tracking environments similar to those of their ancestors under stabilising selection (Ackerly 2009; Losos 2008; Westoby *et al*. 2023). The *separated form* shows how this conservatism is distributed across the network. The climatic effects on *h* and *k_L_* were expressed in conserved variation, so the environmental structuring of stature and hydraulic supply reflects divergence among lineages rather than species-specific deviations. This is consistent with environmental phylogenetic conservatism (Sanchez-Martinez et al. 2024), whereby strongly conserved climatic niches (λPC1 = 0.93) transmit phylogenetic signal to the traits they shape. In contrast, the *k_L_*-*gs* coupling operated at similar strength in both components, and gs inherited most of its phylogenetic signal through this coupling and the edaphoclimatic axes (*λ_eq_* = 0.19 vs. *λ_marg_* = 0.46). Stomatal conductance therefore tracks hydraulic supply regardless of shared ancestry, carrying the phylogenetic imprint reported for stomatal regulation (Brodribb & McAdam 2017; Martin-StPaul *et al*. 2017) without being restricted to it.

Some limitations remain. *k_L_* was compiled from heterogeneous methods and taken as the highest observed value per species and site, so its magnitude depends on sampling effort and method, and sap-flow data concentrated in taller species. The *h*-*k_L_* relationship nevertheless held within methods (Note S2). *k_L_* was not normalised to a reference temperature, so a viscosity-related thermal component may contribute to the climatic effects we report. Finally, *Hv* and *gs* were available for fewer species than *k_L_* and *h*, so paths involving them are estimated with lower precision.

## Conclusion

Across the seed-plant phylogeny, the water supply reaching each unit of leaf area (*k_L_*) declines with plant height. This extends the hydraulic-limitation hypothesis from the growth of individual trees to variation among species. The decline is partly compensated, plausibly by conduit widening towards the stem base and by adjustments in root and leaf xylem, but not by allocation. Taller species do not build more sapwood per unit leaf area, and ultimately operate at both lower *k_L_* and lower concurrent stomatal conductance. The tight coupling between *k_L_*and *gs*, expressed in both conserved and phylogenetically independent variation, indicates that matching stomatal water loss to hydraulic supply is a general and evolutionarily persistent feature of plant design. The climatic effect on *k_L_* was expressed mainly in conserved variation, whereas the height effect could not be attributed to either component. Testing hydraulic limitation within species, using the species × site structure of the data, would clarify how much of the height constraint is inherited and how much reflects plastic or ontogenetic adjustment.

## ACKNOWLEDGMENTS

We thank the authors of the studies compiled in the Kplant database, and the coauthors of the database, without whose published measurements this synthesis would not have been possible. VF would like to thank funding from the Spanish Ministry of Science and Universities, MICIU/AEI/10.13039/501100011033, with the ’Severo Ochoa’ Centres of Excellence programme.

## CONTRIBUTION

VF and MM conceived and designed the study. VF assembled and curated the analysis dataset. LY developed the phylogenetic structural equation model and implemented the path decomposition. VF performed the analyses and wrote the first draft with input from MM and LY. All authors contributed to interpretation and revision, and approved the final manuscript.

---

**Figure S1.**
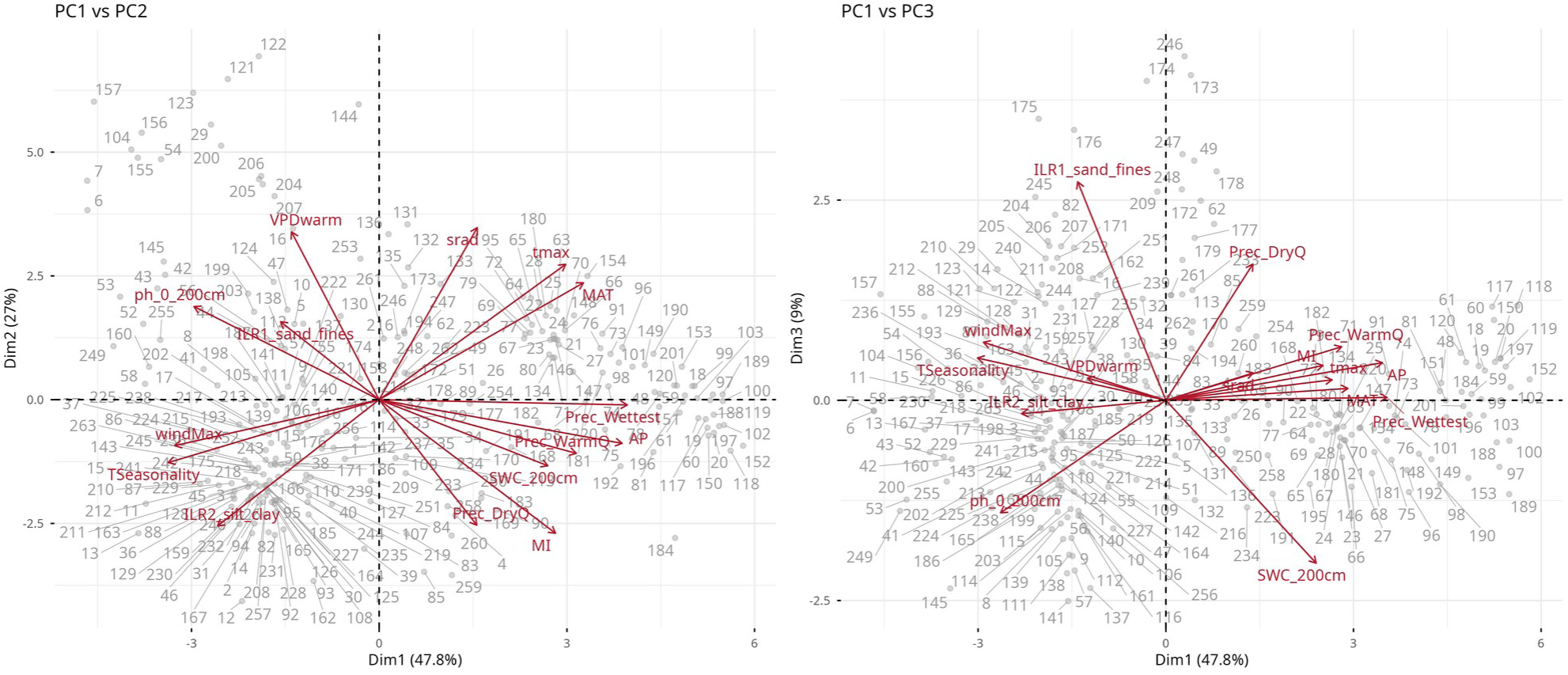
Principal component analysis of the species-level edapho-climatic variables, shown as (a) PC1 versus PC2 and (b) PC1 versus PC3. Arrows show the loadings of each variable; points and values represent different species. PC1 increases toward warm, wet, weakly seasonal niches; PC2 toward high maximum vapour pressure deficit and solar radiation; PC3 toward sandier soils. The first three components together explained 83.8% of the total variance. Variables: MAT, mean annual temperature; tmax, mean annual maximum temperature; TSeasonality, temperature seasonality; AP, annual precipitation; Prec_Wettest, precipitation of the wettest month; Prec_WarmQ, precipitation of the warmest quarter; Prec_DryQ, precipitation of the driest quarter; srad, mean annual solar radiation; windMax, maximum monthly wind speed; VPDwarm, mean vapour pressure deficit of the warmest month; MI, moisture (aridity) index; ph_0_200cm, soil pH (0-200 cm); ILR1_sand_fines, isometric log-ratio of sand relative to silt and clay; ILR2_silt_clay, isometric log-ratio of silt relative to clay; SWC_200cm, volumetric water content at field capacity (−33 kPa) at 100-200 cm

**Figure S2.**
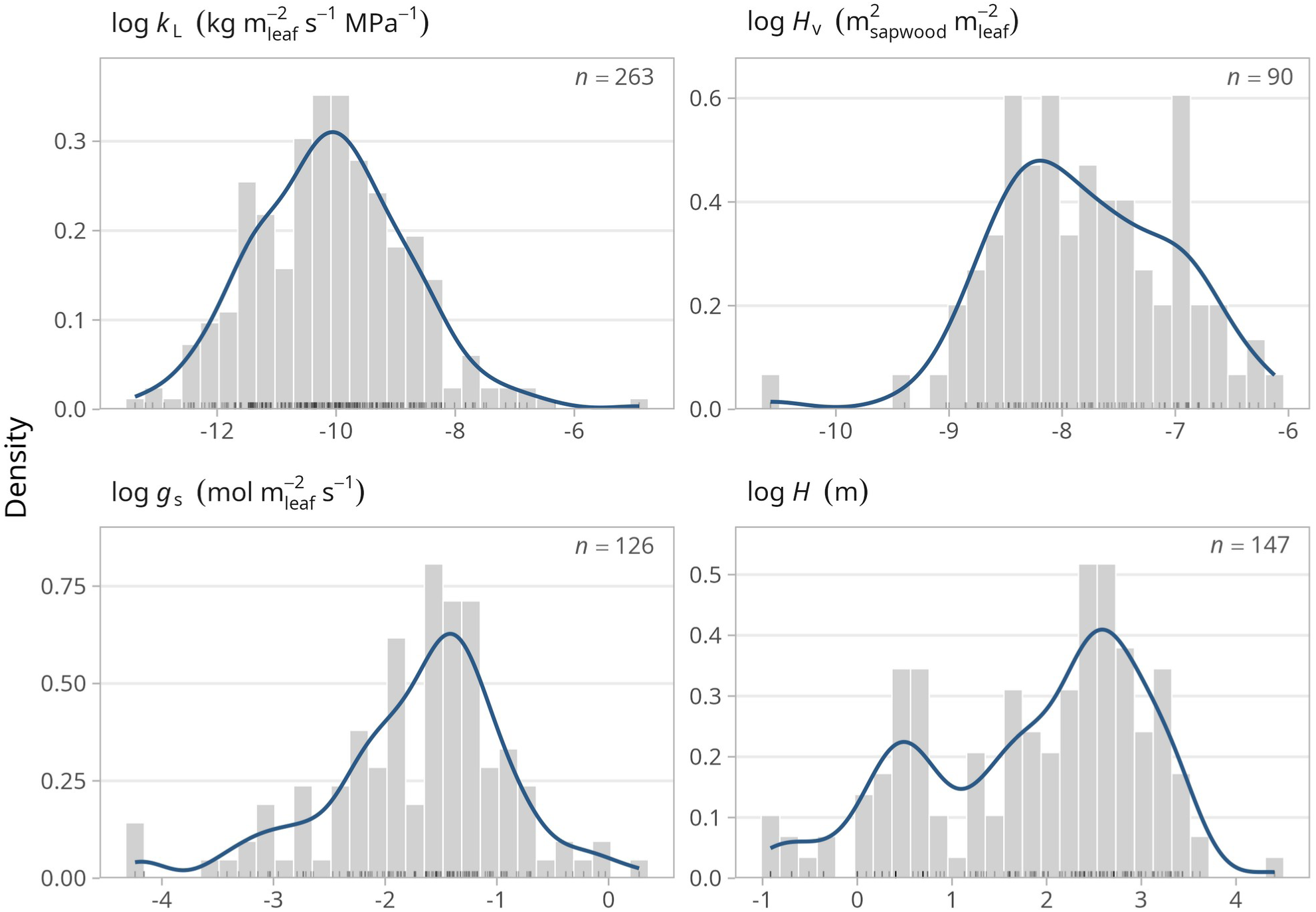
Distributions of the four species-level plant traits after log-transformation: leaf-specific whole-plant hydraulic conductance (*k_L_*), Huber value (*Hv*), stomatal conductance (*gs*) and plant height (*h*).

**Figure S3.**
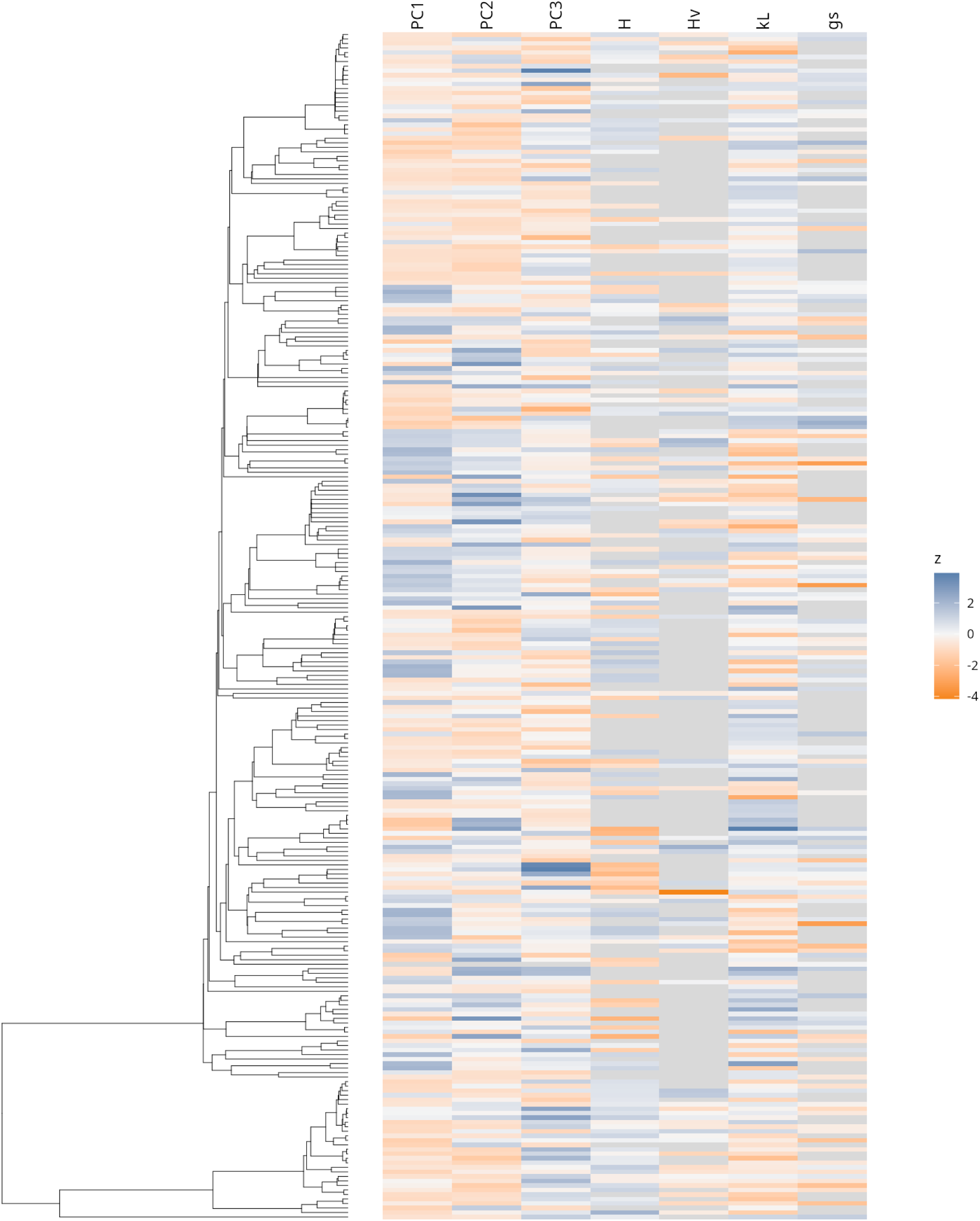
The 263-species tree with the z-scored m7 trait matrix on the tips. Grey marks missing cells.

**Figure S4.**
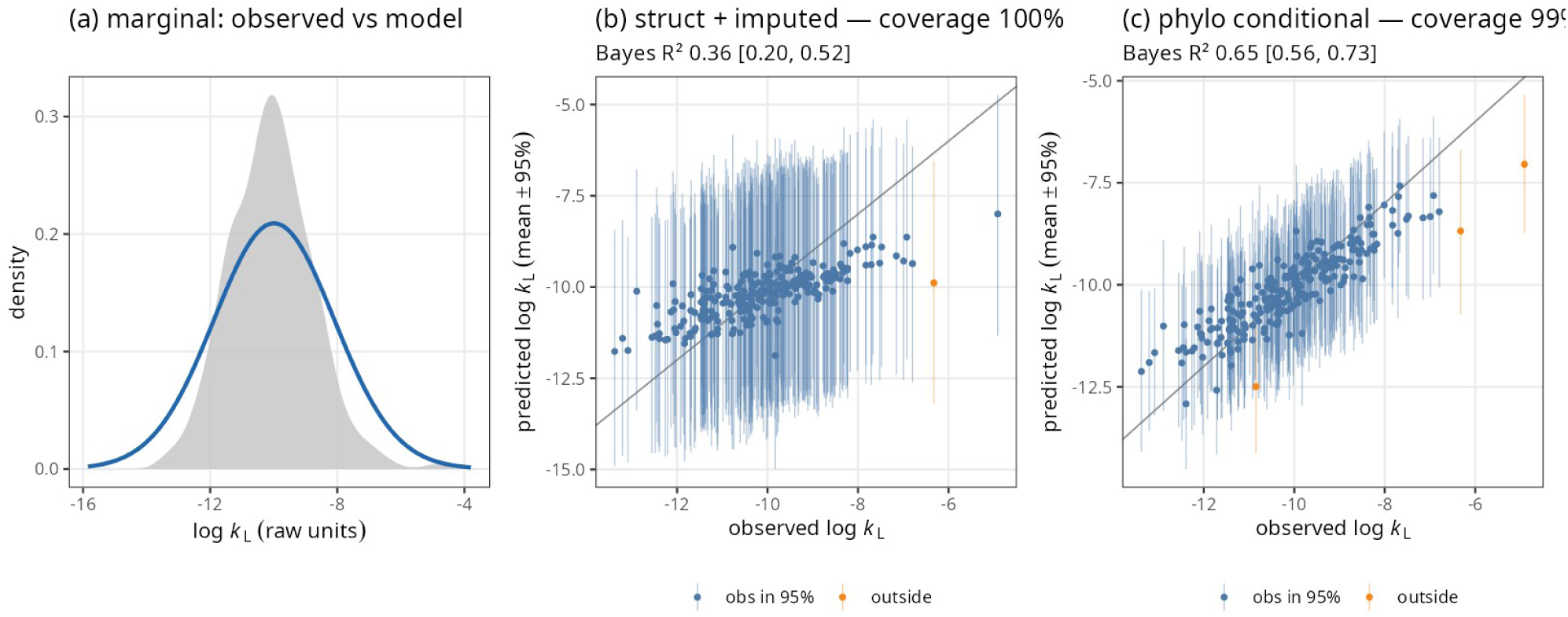
Posterior predictive checks for *k_L_* in the s*hared form* m7 model. (a) Model-implied marginal distribution of *k_L_* (line) against the observed distribution (shaded). (b) Structural prediction: each species’ *k_L_* predicted from its parents in the network, with unmeasured parents filled by their imputed values. (c) Phylogenetic prediction: each species’ *k_L_*predicted conditional on everything else in the dataset, including the *k_L_*of its relatives, through the joint covariance at the tips. Points show posterior means with 95% intervals; the line is 1:1, and orange colour indicates whether the observation falls inside the interval.

**Figure S5.**
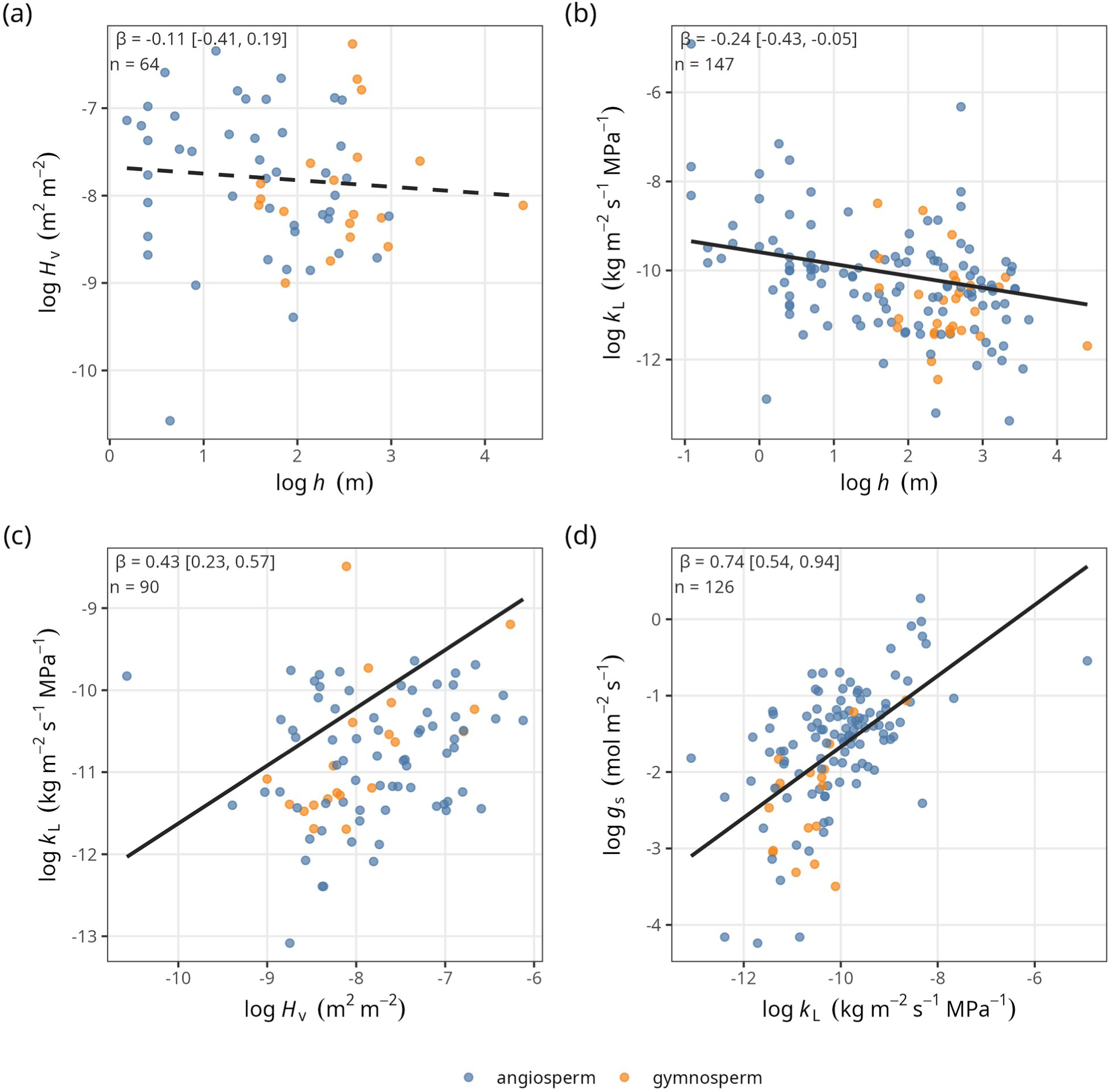
Species-level relationships among traits in the *shared* m7 network: (a) *Hv* against *h*, (b) *k_L_* against *h*, (c) *k_L_* against *Hv*, and (d) *gs* against *k_L_*. Points show observed species values on log scales, coloured by clade (angiosperms, gymnosperms). Lines show the direct effect estimated by the phylogenetic structural equation model, back-transformed to log-log units and drawn through the trait means; because these effects are conditional on the edaphoclimatic axes and the other predictors in the network, they need not match the bivariate trend. Solid lines denote effects whose 95% credible interval excludes zero; dashed lines, effects whose interval overlaps zero. β, standardised direct effect with its 95% credible interval; n, number of species with both variables observed.

**Figure S6.**
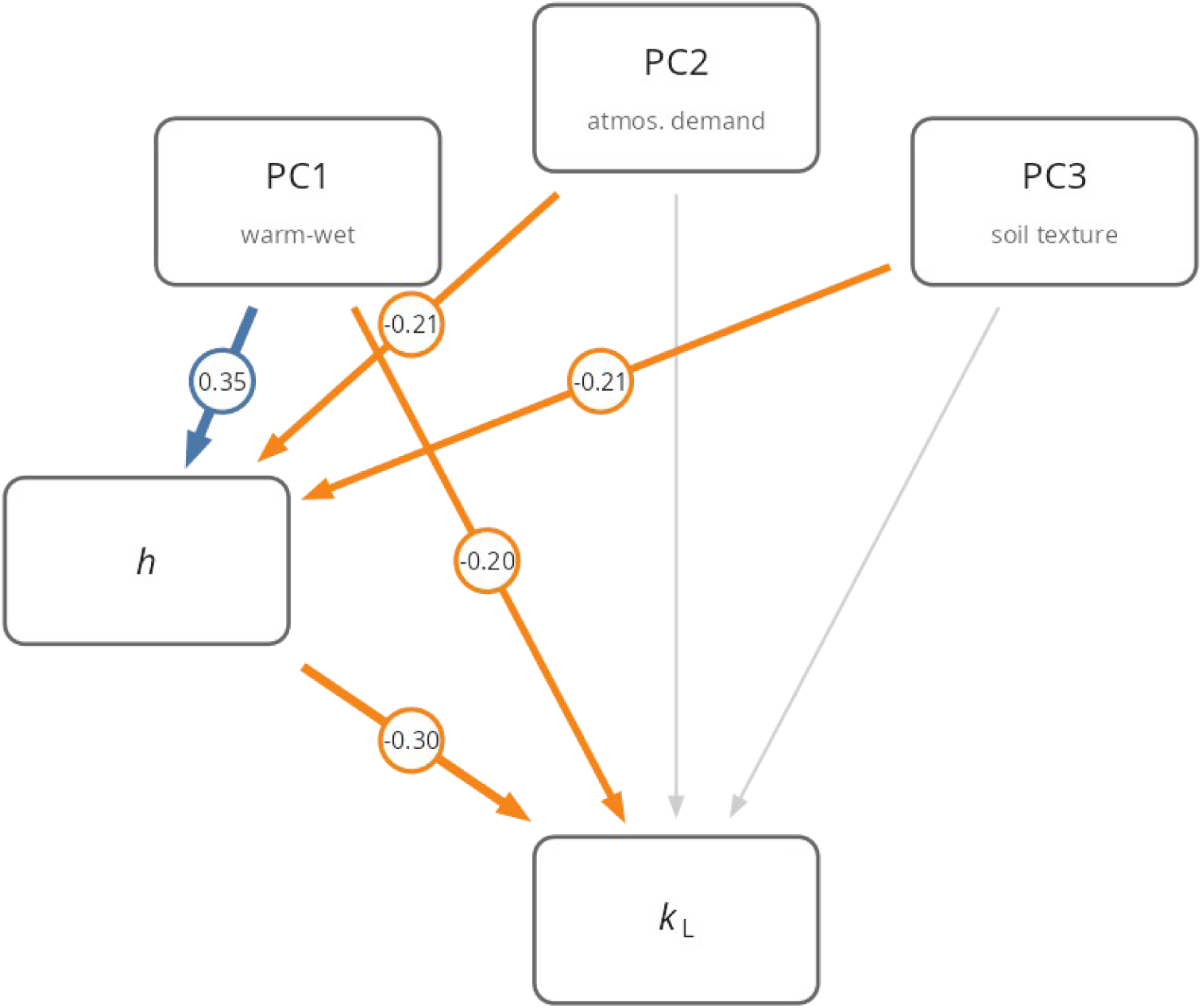
Figure S4. Reduced s*hared form* m5 model showing the paths linking edaphoclimatic axes (PC1-PC3) and height to *k_L_*, fitted to the subset of species with complete observations for all variables (N = 147) and therefore without imputation. Blue arrows denote paths whose 95% credible interval excludes zero and is positive, orange negative paths, and grey paths whose interval overlaps zero; arrow width scales with the absolute standardised coefficient, shown alongside each path. PC1 increases toward warm, wet, weakly seasonal niches; PC2 toward high VPD and solar radiation; PC3 toward sandier soils.

**Figure S7.**
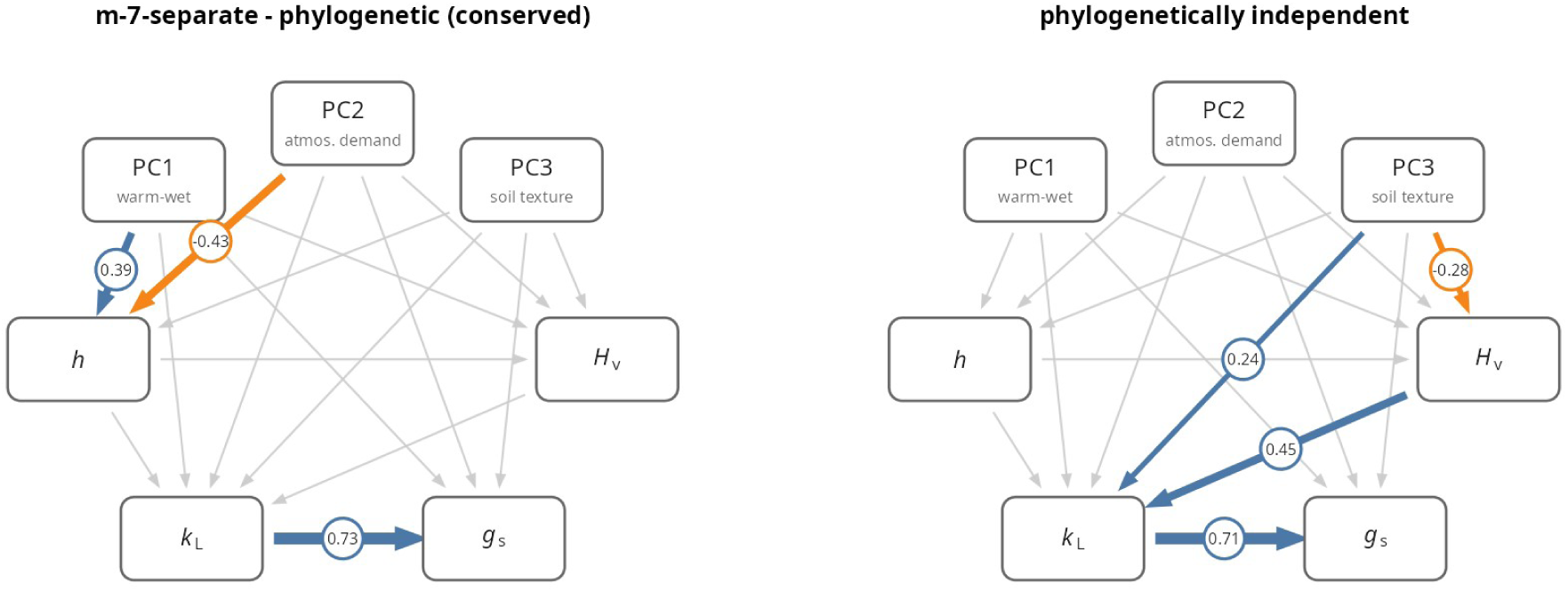
Conserved and phylogenetically independent path networks from the s*eparated form* m7 model, in which the phylogenetically conserved and phylogenetically independent components are estimated independently. (a) Conserved (phylogenetic) network; (b) independent (residual) network. Arrow conventions as in Fig. 2: blue paths have a 95% credible interval excluding zero and positive, orange negative, and grey overlapping zero; arrow width scales with the absolute standardised coefficient, shown alongside each path. PC1 increases toward warm, wet, weakly seasonal niches; PC2 toward high VPD and solar radiation; PC3 toward sandier soils.

**Figure S8.**
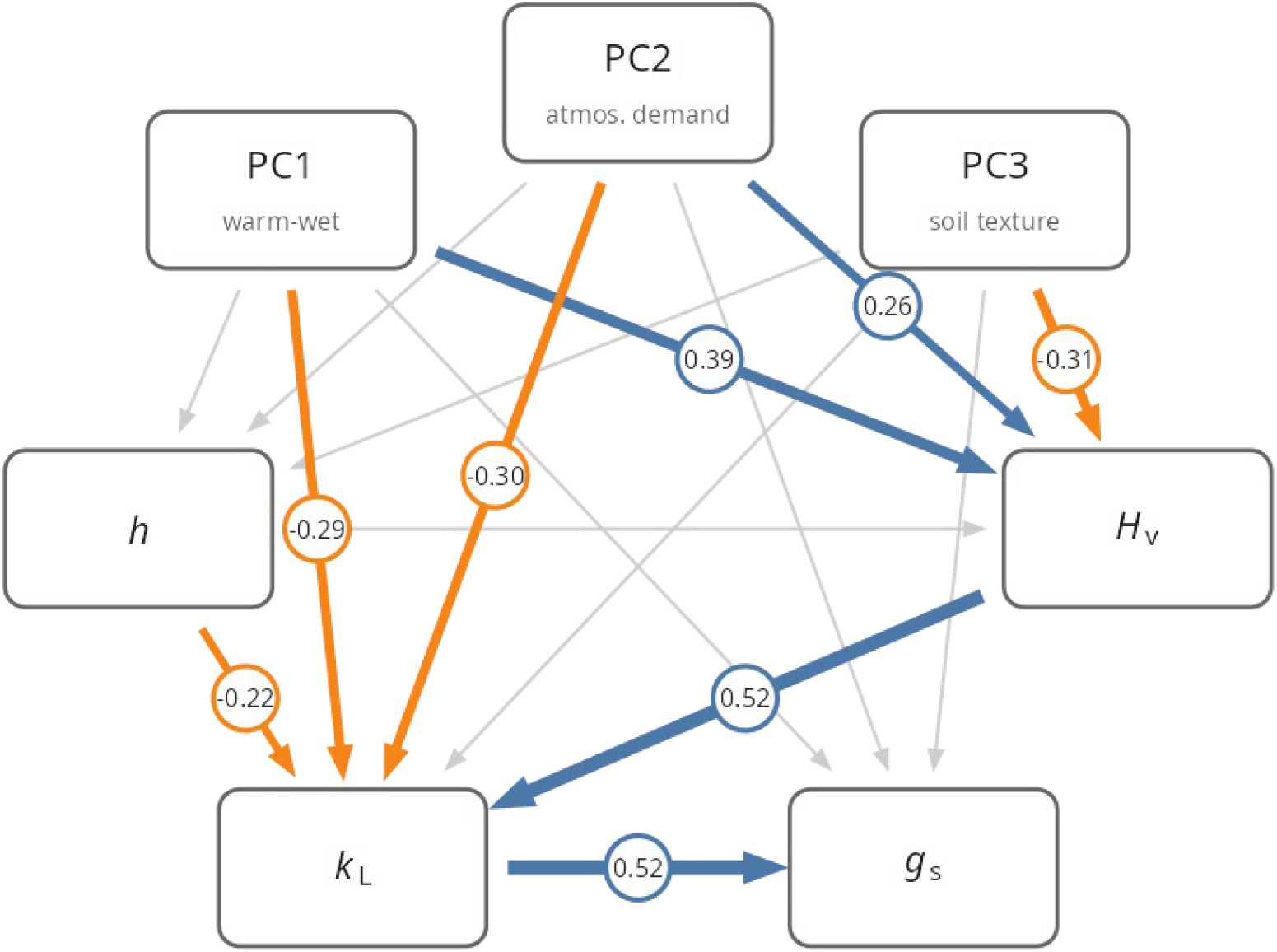
Network of the s*hared form* m7 model fitted to the subset of species whose maximum *k_L_* included sap-flow measurements on one or more individuals (N = 135), showing the direct and indirect paths linking edaphoclimatic axes (PC1-PC3), *h* and *Hv* to *k_L_* and *gs*. Blue arrows denote paths whose 95% credible interval excludes zero and is positive, orange negative paths, and grey paths whose interval overlaps zero; arrow width scales with the absolute standardised coefficient, shown alongside each path. PC1 increases toward warm, wet, weakly seasonal niches; PC2 toward high VPD and solar radiation; PC3 toward sandier soils.

**Figure S9.**
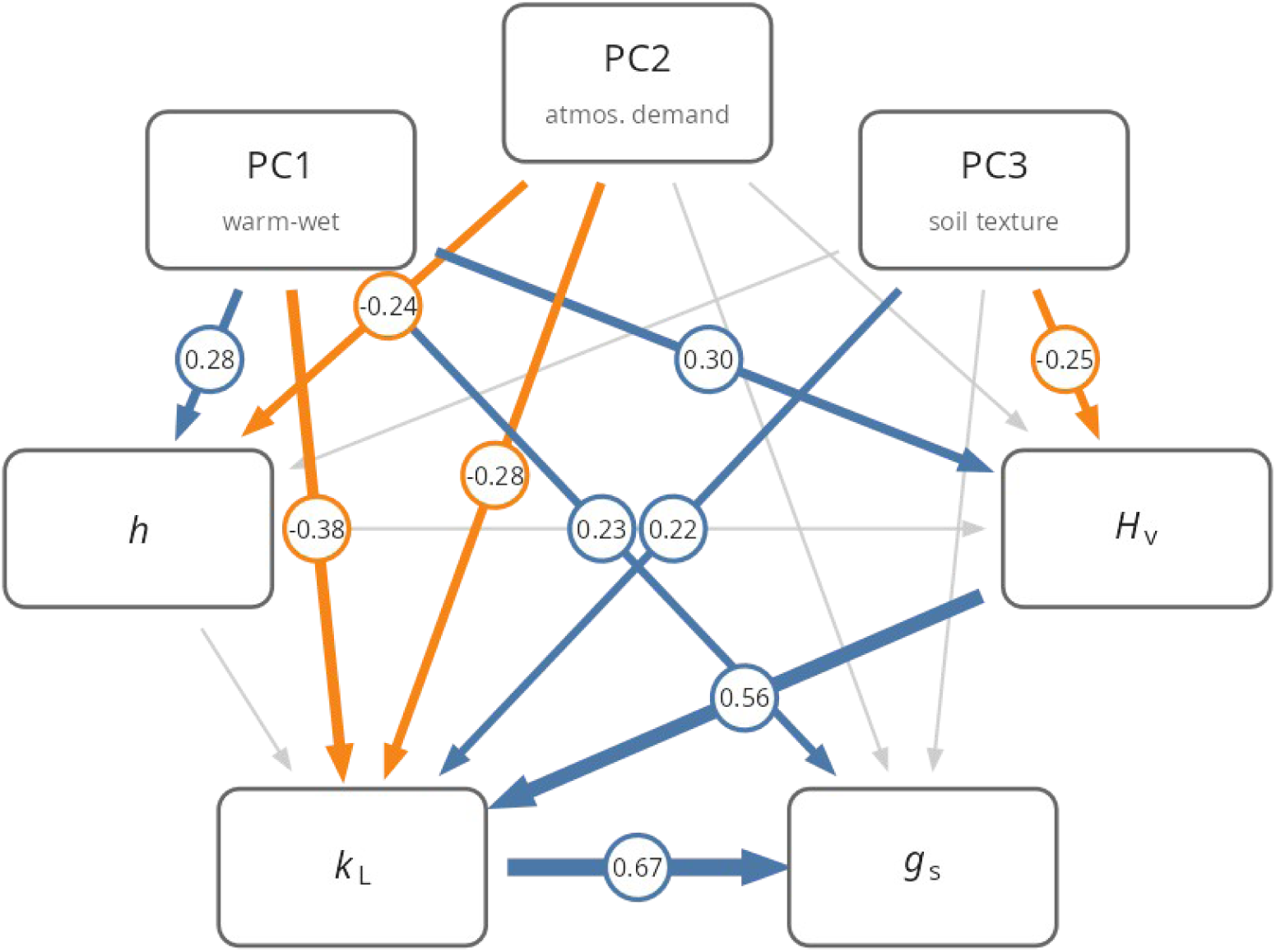
Network of the s*hared form* m7 model fitted to trees only (N = 192), showing the direct and indirect paths linking edaphoclimatic axes (PC1-PC3), *h* and *Hv* to *k_L_* and *gs*. Blue arrows denote paths whose 95% credible interval excludes zero and is positive, orange negative paths, and grey paths whose interval overlaps zero; arrow width scales with the absolute standardised coefficient, shown alongside each path. PC1 increases toward warm, wet, weakly seasonal niches; PC2 toward high VPD and solar radiation; PC3 toward sandier soils.

**Figure S10.**
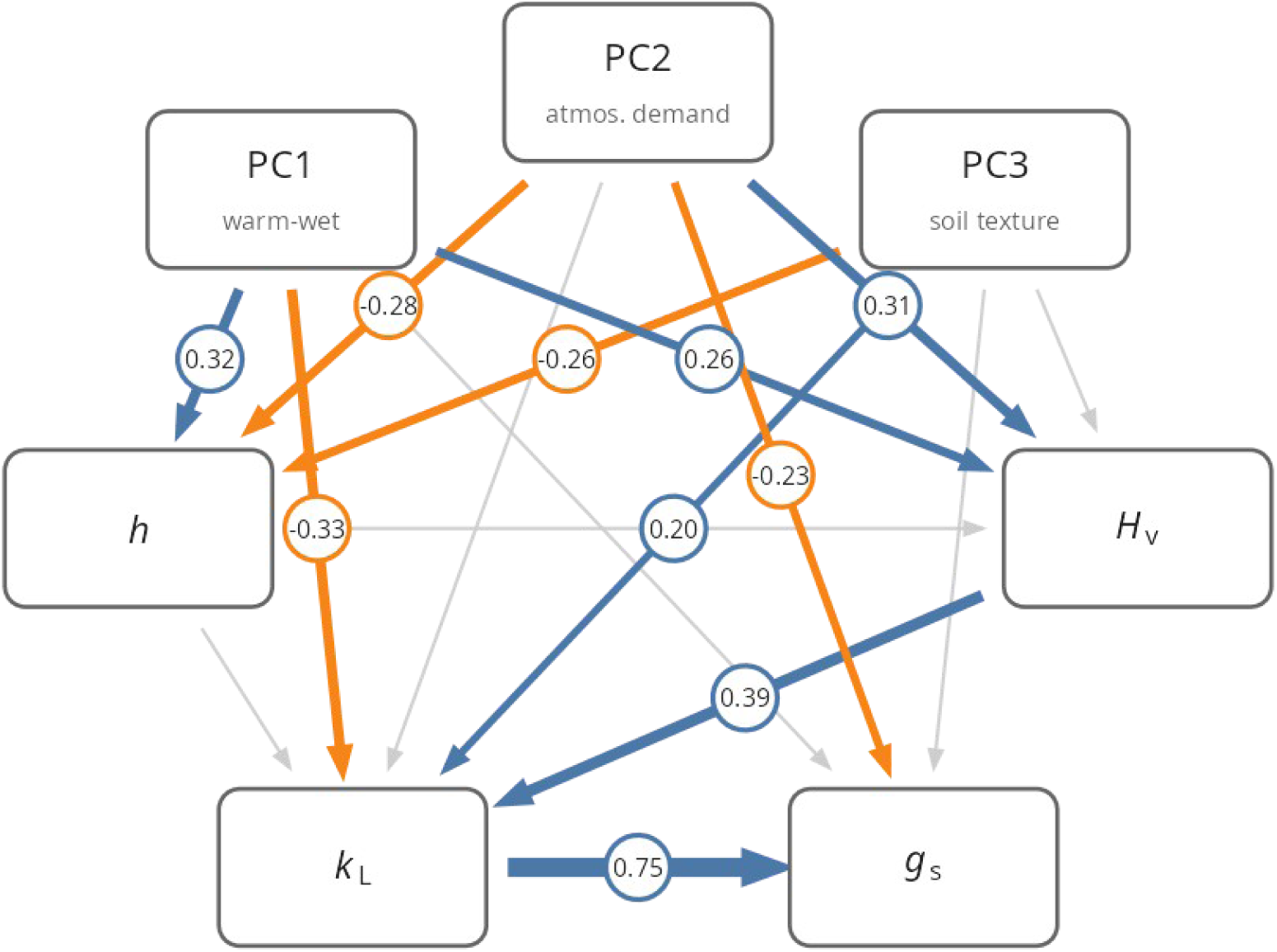
Network of the s*hared form* m7 model fitted to angiosperm species only (N = 232), showing the direct and indirect paths linking edaphoclimatic axes (PC1-PC3), *h* and *Hv* to *k_L_* and *gs*. Blue arrows denote paths whose 95% credible interval excludes zero and is positive, orange negative paths, and grey paths whose interval overlaps zero; arrow width scales with the absolute standardised coefficient, shown alongside each path. PC1 increases toward warm, wet, weakly seasonal niches; PC2 toward high VPD and solar radiation; PC3 toward sandier soils.

**Table S1.** Treatment and condition levels retained in the analysis, grouped by descriptor family. From the source studies compiled in the Kplant database (Zhao et al. 2026), we retained only records corresponding to field-grown, untreated adult plants measured under non-limiting conditions; the levels listed represent the accepted “si_treatment” categories after this filtering, organised by the type of condition they describe. Different treatments are separated by “/”.

| **Category** | **Treatment and condition levels** |
|---|---|
| Water regime | 0K/+W / 100irrigationneeds / 30% irrigation / 45irrigationneeds / 60% irrigation / Afternoon irrigation (A) / Control (natural water regime) / Deep irrigation (once a week) / Irrigation / Morning irrigation (M) / Post Irrigation / RPI / RainFed / Recovery / dry / heteregenous soil water conditions; control (irrigated) / homogeneous soil water conditions; control (irrigated) / irrigated (IR) / irrigated vines / irrigation / natural variation in soil moisture / non-irrigated (ST) / non-irrigated vines / wet |
| Groundwater / soil depth | B2. 50 m distance from river 4.91m Groundwater depth / B3. 100 m distance from river 6.45m Groundwater depth / E4. 550 m distance from river 6.36 m Groundwater depth / E5. 850m distance from river 8.44m Groundwater depth / deep groundwater / shallow groundwater / soil depth |
| Atmospheric CO2 / VPD | Ambient CO2 / Ambient CO2 1KPa VPD / Ambient CO2 3KPa VPD / Ambient atmospheric CO2 / atmospheric CO2_lower crown / atmospheric CO2_upper crown |
| Canopy / crown position | Control - Lower crown - Whole plant / Control - Upper crown - Whole plant / Intact branches in the lower canopy / Lower canopy; above psi_opt / Lower canopy; bellow psi_opt / Lower crown / Upper canopy; above psi_opt / Upper canopy; bellow psi_opt / Upper crown / bottom canopy / intact branches in the upper canopy / lower canopy / top canopy / upper canopy |
| Season / phenology | DRY season / Early summer / Late summer / WET season / Wet season / control - dry season / control - wet season / dry season / early season / late season / mid-season / spring / summer / wet season |
| Light environment | shade / shade-grown / sun |
| Habitat / site | Bordeaux / Fundao / Lowland / Native range / Park / Plantation / Pond / Street / Upland / bog-grown / forest / hedgerow / restored / savanna |
| Age / shoot type | 1 year old resprout from 16-year-old shrubs / Reiterated shoots / Sequential shoots / adults |
| Management / health | non-infected / ungrazed |
| Rootstock | Rootstock: Actinidia hemsleyana / Rootstock: Actinidia kolomikta / Rootstock: Actinidia macrosperma / Rootstock: Actinidia polygama / Rootstock: CG202 / Rootstock: CG30 / Rootstock: CG934 / Rootstock: M111 / Rootstock: M7A / Rootstock: M9 / Rootstock: MM106 / grown on a rootstock of P. domestica |
| Size / plot | 10 m plot / 15 m / 25 m plot / 30 m / 60 m / intermediate / large / short / tall / tree height |
| Control / ambient | Ambient / CONTROL / Control / contro / control |

**Table S2.** Phylogenetic signal per trait (*λ_eq_* and *λ_marg_*, as in Table 2) from the s*hared form* m7 model fitted to each subset: species whose maximum *k_L_* included sap-flow-based estimates (N = 135), trees only (N = 192), and angiosperms only (N = 232). Values are posterior means with 95% credible intervals.

| Trait | Sap flow *λ<sub>eq</sub>* | Sap flow *λ<sub>marg</sub>* | Trees λ<sub>eq</sub> | Trees *λ<sub>marg</sub>* | Angiosperms *λ<sub>eq</sub>* | Angiosperms *λ<sub>marg</sub>* |
|---|---|---|---|---|---|---|
| PC1 | 0.95 [0.90, 0.99] | 0.95 [0.90, 0.99] | 0.95 [0.90, 0.98] | 0.95 [0.90, 0.98] | 0.86 [0.76, 0.93] | 0.86 [0.76, 0.93] |
| PC2 | 0.68 [0.42, 0.87] | 0.68 [0.42, 0.87] | 0.78 [0.62, 0.89] | 0.78 [0.62, 0.89] | 0.65 [0.47, 0.79] | 0.65 [0.47, 0.79] |
| PC3 | 0.34 [0.08, 0.62] | 0.34 [0.08, 0.62] | 0.13 [0.01, 0.37] | 0.13 [0.01, 0.37] | 0.14 [0.00, 0.38] | 0.14 [0.00, 0.38] |
| *h* | 0.77 [0.42, 0.94] | 0.78 [0.47, 0.93] | 0.33 [0.01, 0.76] | 0.47 [0.14, 0.79] | 0.43 [0.16, 0.68] | 0.50 [0.30, 0.68] |
| *Hv* | 0.40 [0.01, 0.83] | 0.56 [0.25, 0.83] | 0.21 [0.00, 0.65] | 0.36 [0.10, 0.67] | 0.19 [0.00, 0.57] | 0.29 [0.09, 0.58] |
| *k<sub>L</sub>* | 0.46 [0.01, 0.84] | 0.54 [0.21, 0.79] | 0.34 [0.06, 0.67] | 0.43 [0.23, 0.63] | 0.48 [0.12, 0.75] | 0.48 [0.29, 0.65] |
| *gs* | 0.21 [0.00, 0.57] | 0.40 [0.18, 0.63] | 0.21 [0.00, 0.54] | 0.34 [0.15, 0.55] | 0.08 [0.00, 0.32] | 0.27 [0.13, 0.43] |

**Table S3.**
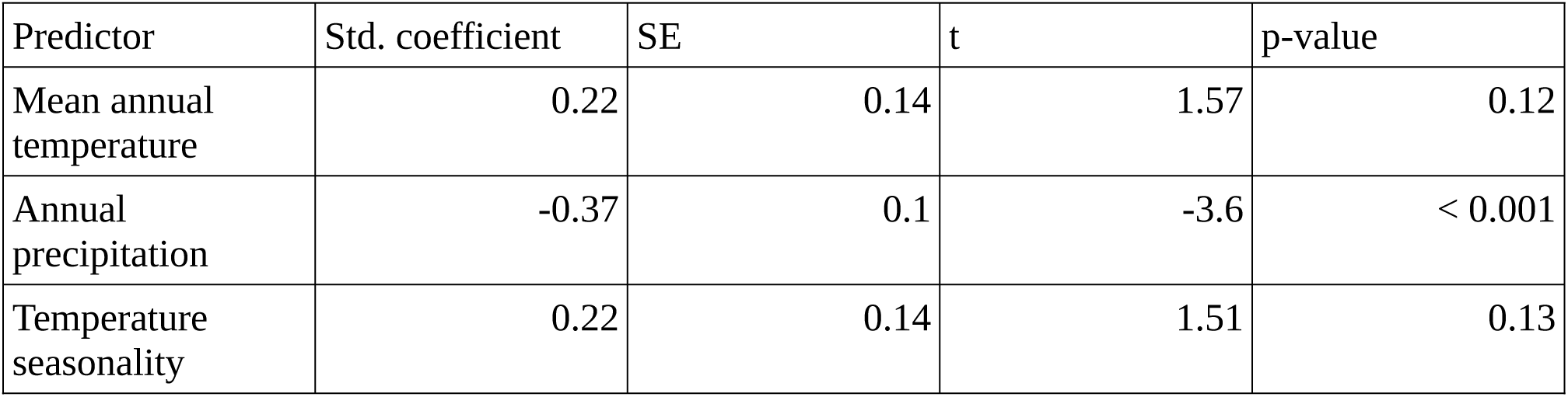
Direct associations between climate and leaf-specific whole-plant conductance (*k_L_*). Coefficients are from an ordinary least-squares regression of species-mean log(*k_L_*) on mean annual temperature, annual precipitation and temperature seasonality, with predictors standardised to zero mean and unit variance so that coefficients are comparable in magnitude (N = 263; model RZ = 0.096).

| Predictor | Std. coefficient | SE | t | p-value |
|---|---|---|---|---|
| Mean annual temperature | 0.22 | 0.14 | 1.57 | 0.12 |
| Annual precipitation | -0.37 | 0.1 | -3.6 | < 0.001 |
| Temperature seasonality | 0.22 | 0.14 | 1.51 | 0.13 |

**Table S4.** Decomposition of the effects of the edaphoclimatic axes on *k*_L_ into direct and indirect routes in the *shared* m7 network. Each indirect route is the product of the path coefficients along it, computed for every posterior draw; values are posterior means with 95% credible intervals. The total effect equals the sum of the direct effect and all indirect routes. PC1 increases toward warm, wet niches with low temperature seasonality; PC2 toward high atmospheric demand and radiation; PC3 toward sandier soils with lower deep-soil water retention. * denotes credible interval do not cross zero.

| Predictor | Route | Effect [95% CI] |
|---|---|---|
| PC1 | Direct | -0.31 [-0.46, -0.15]* |
| PC1 | PC1 → Hv → kL | 0.11 [0.01, 0.22]* |
| PC1 | PC1 → h → kL | -0.08 [-0.17, -0.01]* |
| PC1 | PC1 → h → Hv → kL | -0.02 [-0.07, 0.03] |
| PC1 | Total | -0.30 [-0.43, -0.17]* |
| PC2 | Direct | -0.15 [-0.30, -0.00]* |
| PC2 | PC2 → Hv → kL | 0.14 [0.04, 0.26]* |
| PC2 | PC2 → h → kL | 0.05 [0.00, 0.11]* |
| PC2 | PC2 → h → Hv → kL | 0.01 [-0.02, 0.05] |
| PC2 | Total | 0.05 [-0.08, 0.18] |
| PC3 | Direct | 0.21 [0.07, 0.35]* |
| PC3 | PC3 → Hv → kL | -0.12 [-0.24, -0.02]* |
| PC3 | PC3 → h → kL | 0.05 [0.01, 0.11]* |
| PC3 | PC3 → h → Hv → kL | 0.01 [-0.02, 0.04] |
| PC3 | Total | 0.14 [0.03, 0.25]* |

**Table S5.** Sensitivity of key standardised effects to data subset and model structure ( *shared form*). Values are posterior means with 95% credible intervals; asterisks mark intervals excluding zero. “PC1 → *h* → *k_L_*” is the indirect route through h. TRY maximum height: measured *h* replaced by the 95th percentile of TRY height records. Complete cases (m5): species with no missing values. m5 excludes *Hv* and *gs* (-), so its direct environmental effects on *k_L_* are not conditional on *Hv*. Standardisation and tree rescaling were recomputed within each subset.

| effect | All species (m7) | Sap flow | Trees | Angiosperms | TRY maximum height | All species (m5) | Complete cases (m5) |
|---|---|---|---|---|---|---|---|
| N species | 263 | 135 | 192 | 232 | 263 | 263 | 147 |
| h → kL (direct) | -0.24 [-0.43, -0.05]* | -0.22 [-0.42, -0.02]* | -0.13 [-0.30, 0.04] | -0.22 [-0.43, 0.00] | -0.18 [-0.36, 0.01] | -0.30 [-0.47, -0.11]* | -0.31 [-0.50, -0.11]* |
| h → kL (total) | -0.29 [-0.47, -0.10]* | -0.18 [-0.39, 0.05] | -0.20 [-0.37, -0.02]* | -0.28 [-0.48, -0.07]* | -0.29 [-0.43, -0.14]* | -0.30 [-0.47, -0.11]* | -0.31 [-0.50, -0.11]* |
| h → Hv | -0.11 [-0.41, 0.19] | 0.08 [-0.20, 0.36] | -0.12 [-0.38, 0.14] | -0.16 [-0.52, 0.20] | -0.25 [-0.55, 0.04] | NA | NA |
| Hv → kL | 0.43 [0.23, 0.57]* | 0.52 [0.29, 0.70]* | 0.56 [0.39, 0.71]* | 0.39 [0.12, 0.57]* | 0.44 [0.26, 0.58]* | NA | NA |
| kL → gs | 0.74 [0.54, 0.94]* | 0.52 [0.22, 0.81]* | 0.67 [0.44, 0.89]* | 0.75 [0.52, 0.97]* | 0.74 [0.53, 0.93]* | NA | NA |
| PC1 → kL (direct) | -0.31 [-0.46, -0.15]* | -0.29 [-0.51, -0.05]* | -0.38 [-0.55, -0.21]* | -0.33 [-0.50, -0.16]* | -0.33 [-0.47, -0.19]* | -0.20 [-0.34, -0.05]* | -0.03 [-0.22, 0.16] |
| PC1 → h → kL | -0.08 [-0.17, -0.01]* | -0.05 [-0.14, 0.00] | -0.04 [-0.11, 0.01] | -0.07 [-0.16, 0.00] | -0.04 [-0.09, 0.00] | -0.10 [-0.20, -0.03]* | -0.10 [-0.19, -0.03]* |
| PC1 → kL (total) | -0.30 [-0.43, -0.17]* | -0.14 [-0.35, 0.09] | -0.27 [-0.43, -0.11]* | -0.32 [-0.46, -0.17]* | -0.29 [-0.42, -0.16]* | -0.30 [-0.43, -0.17]* | -0.13 [-0.31, 0.06] |
| PC2 → kL (direct) | -0.15 [-0.30, -0.00]* | -0.30 [-0.48, -0.11]* | -0.28 [-0.43, -0.14]* | -0.16 [-0.33, 0.00] | -0.13 [-0.27, 0.01] | -0.01 [-0.15, 0.12] | 0.13 [-0.04, 0.31] |
| PC2 → kL (total) | 0.05 [-0.08, 0.18] | -0.14 [-0.33, 0.06] | -0.14 [-0.30, 0.01] | 0.04 [-0.10, 0.17] | 0.05 [-0.07, 0.18] | 0.05 [-0.08, 0.18] | 0.21 [0.03, 0.38]* |
| PC3 → kL (direct) | 0.21 [0.07, 0.35]* | 0.08 [-0.10, 0.26] | 0.22 [0.07, 0.37]* | 0.20 [0.04, 0.37]* | 0.25 [0.12, 0.38]* | 0.08 [-0.03, 0.20] | 0.05 [-0.11, 0.22] |
| PC3 → kL (total) | 0.14 [0.03, 0.25]* | -0.08 [-0.26, 0.09] | 0.11 [-0.03, 0.24] | 0.16 [0.04, 0.28]* | 0.15 [0.04, 0.26]* | 0.15 [0.04, 0.25]* | 0.12 [-0.05, 0.28] |

## Note S1. Model specification, estimation and derived quantities

### S1.1 Structure

Each species’ K-vector of traits (or edaphoclimatic variables) was modelled as

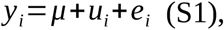

where *i*=1 , … *, N* indexes species, *μ* is the vector of ancestral means, *ui* the phylogenetically conserved component and *ei* the phylogenetically independent component.

Within each component c ∈ {phy, ind} the traits follow a recursive path model in causal order, so the component covariance takes the structural form

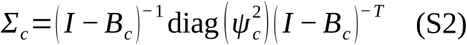

with *Bc* strictly lower-triangular and non-zero only on the retained directed acyclic graph edges; dropped edges are fixed to zero rather than estimated parameters. This class of path-constrained phylogenetic models are a sub-class of multi-response phylogenetic mixed models which are typically used to estimate unconstrained covariance matrices for each component (Halliwell *et al*. 2025).

Two path structures are fitted:

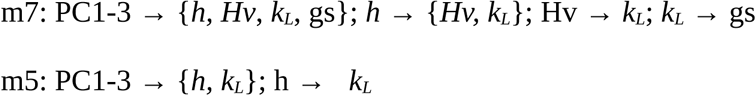

Each of these is used in two model forms. In the s*hared form*, one coefficient matrix B enters both components, so the conserved, phylogenetically independent and phenotypic-total networks coincide, sharing the same estimated path coefficients and only the disturbance variances differ between components. In the s*eparated form*, B_phy_ and B_ind_ are estimated separately, which allows a conserved and a phylogenetically independent path to differ in both magnitude and sign.

### S1.2 Likelihood and eigen-rotation

Index tips i = 1, …, N and traits k = 1, …, K in causal order. Collect the traits in the N × K matrix Y = (y_ik), where i indexes species (rows) and k indexes traits (columns). Missing values are estimated as model parameters and inserted into *Y* per draw (S1.3). Marginalising the Brownian process over the tree, the tips follow a single multivariate normal,

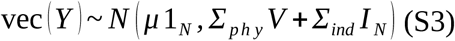

where vec stacks Y by column and V is the tip correlation matrix implied by Brownian motion, that is, the phylogenetic variance-covariance matrix of the tree rescaled to unit root-to-tip height.

To reduce the computational cost of fitting the model, we eigendecompose *V* =*UΛ U^T^*once before fitting and use the orthogonal U to rotate the species (rows) *Y* *=*U^T^ Y* which diagonalises the phylogenetic term while leaving the independent one unchanged (*U^⊤^ U* =*I* ) ,. This transformation preserves the likelihood and factorises the NK-variate normal into N independent K-variate normals such that each rotated row a has mean (*U^T^* 1 )_and covariance *^ν^a· ^Σ^ph y* ^+ *Σ*^*ind*, where *^ν^a* is the *a^t^ ^h^* eigenvalue of *V* .

### S1.3 Missing data

Missing trait measurements are sampled as parameters, so the rotation in S1.2 acts on complete data. The missing values are imputed at every MCMC iteration jointly with the remaining parameters, conditional on the phylogeny and on the trait network.

### S1.4 Priors and estimation

Writing the component coefficient matrix as *Bc* =(*bc , kl* ), see Equations (S2) and (S3), the priors for the model are

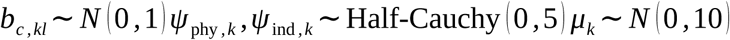

In the s*hared form* case, one set of *bkl* serves both components.. Traits are z-scored and the tree rescaled to unit root-to-tip height. All model variants are specified in Stan programming language (Carpenter *et al*. 2017) and fitted with cmdstanr (CmdStan 2.38), using four chains of 4000 iterations (2000 warm-up), adapt_delta = 0.95, and maximum tree depth 12. Models were check for convergence by observation of trace plots and the requirement that all R^S^ values were less than 1.01 and effective sample sizes exceed 400 (Vehtari *et al*. 2021).

### S1.5 Phylogenetic signal

In the SEM context, there are two natural definitions of per-trait phylogenetic signal. The first is the proportion of phylogenetic variation in the disturbance variances of individual trait equations. This estimates the phylogenetic signal in the trait itself ignoring any signal that arises via the parent traits:

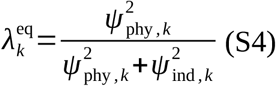

The second is the familiar marginal ratio based on the phenotypic variance decomposition

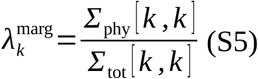

The two forms are equivalent if there are no parent traits (i.e., the exogenous climate PCs).

### S1.6 Effect decomposition

The direct effects correspond to a single arrow in the path diagram and are the entries of the coefficient matrix B. The total effects accumulate every directed route from one trait to another which can be computed for all trait pairs by summing the matrix powers of B or equivalently as ( *I −B* )*^−^*^1^ *− I* (Bollen 1987). The indirect effects are the difference between the total and the direct, ( *I − B* )*^−^*^1^ *− I − B*. All three sets of effects are evaluated on each posterior draw yielding a posterior distribution for each, summarised by their mean and 2.5th and 97.5th percentiles.

### S1.7 Predictive checks for *k_L_*

To test the ability of the model to generate plausible data, two posterior predictive checks are shown in Fig. S4. The first predicts each species’ *k_L_* from its parents in the network, filling in any unmeasured parent with its imputed value. The second predicts a single species’ *k_L_* value conditional on everything else in the dataset, including the *k_L_* of its relatives, exploiting the joint covariance at the tips. The first check is the more difficult prediction problem since it relies solely on the fitted network and the observed traits for each species. The second check tests predominantly the degree to which phylogenetic signal in the response can explain the observed variation.

## Note S2. Allometric scaling of *k_L_* with height

We linearly regressed species-mean log leaf-specific whole-plant conductance on log height (*h*): log *k_L_* = β₀ + β₁ log *h* The slope β₁ is the allometric exponent of *k_L_*∝ *h*^β₁. Fitted on angiosperms (n = 119): β₁ = -0.42 (SE 0.086, p = 3×10^-6^, RZ = 0.17); and on the full dataset (angiosperms + gymnosperms, n = 147): β₁ = -0.44 (SE 0.078, p = 6×10^-8^, RZ = 0.18). Adding a term for whether the estimate derived from sap flow shifts the intercept but not the slope (interaction p = 0.19), and leaves the exponent well above -1 (-0.32, SE 0.085).

### S2.1 Theoretical framework

Following Magnani et al. (2000; see Mencuccini 2003), whole-plant conductance ( *k_P_*) can be written as

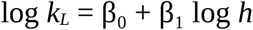

with *Ks* the sapwood-specific conductivity, M_S_ the plant sapwood mass, *h* height and ρs sapwood density. Writing the fractional leaf mass F such that M_L_ = F·M_S_/(1-F), and M_L_ = *A_L_/SLA* (*A_L_*, leaf area; *SLA*, specific leaf area), gives Ms = *A_L_*·(1-F)/(*SLA*·F) and hence, per unit leaf area,

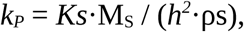

If *Ks*, F, *ρs* and *SLA* were independent of height, *k_L_* would scale as *h*^-2^ (β = -2). Expressing allocation through the Huber value (*Hv* = A_S_/A_L_) makes the geometry explicit: since (1-F)/F = *Hv*·*h*·*ρs*·*SLA*, the same relation can be written as a tube approximation,

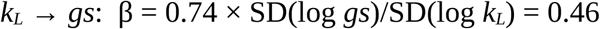

in which local tissue conductivity (*Ks_local*), sapwood-to-leaf allocation and path length appear as three separate terms. Under this approximation, and with *Ks_local* and *Hv* independent of height, *k_L_* instead scales as *h*^-1^ (β = -1).

### S2.2 Why the exponent departs from -1

Our slope (≈ -0.4), like the -0.47 of Mencuccini (2003), lies well above -1, so one or more terms must covary with height. Allocation does not: *Hv* was unrelated to height among species, contributing no compensation. The principal term is conductivity. The tube approximation treats *Ks* as a local, height-independent value, whereas the conductance of the whole path is the integral of conductivity along it, and tip-to-base conduit widening makes the effective, path-integrated conductivity increase with *h* (Olson et al. 2014; Anfodillo & Olson 2024). This positive covariation offsets part of the -1 exponent, while ρs and SLA define further axes along which *k_L_* may vary independently of height.

Equivalently, the exact identity *k_L_* = *Ks_eff*·*Hv* (with *Ks_eff* = (∫₀ʰ dx/Ks(x))⁻¹ the path-integrated conductance per unit sapwood area, which reduces to Ks/h for uniform conductivity, equivalent to k_S_ in Note S2.3) absorbs the height dependence into *Ks_eff*, which is why *h* does not appear as a separate term once conductivity is integrated rather than treated as local. The tube approximation makes the two possible routes to compensation explicit: invariance of *k_L_* (an exponent of zero) requires the numerator *Ks_local*·*Hv* to scale as *h*^1, achievable either by sapwood allocation increasing with height (*Hv* ∝ *h*) or by tissue conductivity doing so (*Ks* ∝ *h*). Our data exclude the first among species, since *Hv* was unrelated to height, leaving conductivity as the only available route. This is the route formalised by tip-to-base conduit widening, through which effective conductivity rises with height and, if sufficient, holds *k_L_* constant (Anfodillo & Olson 2024). That we observe an exponent of ≈ -0.4 rather than zero indicates that this compensation, operating through conductivity alone, is substantial but incomplete.

### S2.3 Allocation, conductance per unit sapwood area and climate

Because *k_L_* = *k_S_*·*Hv* is an identity, with *k_S_* the whole-plant conductance per unit sapwood area, the two quantities are related in logs by log *k_L_* = log *k_S_* + log *Hv.* Regressing either side on log *Hv* therefore gives slopes differing by exactly one. Across the 90 species with an observed *Hv*: log *k_L_* on log *Hv*: β = 0.25 (SE 0.10, p = 0.017) log *k_S_* on log *Hv*: β = -0.75 (SE 0.10, p = 1×10^-10^)

To identify which climatic component of PC1 underlies its positive effect on *Hv*, we also regressed log *Hv* on mean annual temperature (MAT) and annual precipitation (AP) across the same 90 species. *Hv* increased with MAT (0.044 °C^-1^, SE 0.014, p = 0.002; R^2^ = 0.12) but was unrelated to AP (p = 0.70).

### S2.4 Relation to the SEM coefficients

The *h* → *k_L_* path coefficient in the phylogenetic SEM is not directly comparable to β₁: it is standardised (traits z-scored) and conditional on the other predictors (climate axes and *Hv*). The standardised coefficient from the *shared form* m7 (-0.24) back-transforms to original units as β = -0.24 × SD(log *k_L_*)/SD(log *h*) = -0.27, the conditional allometric slope, comparable to β₁ in units but not in interpretation, since it is the effect of height net of the other paths.

Same applies for the rest of the paths:

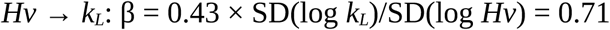

### S2.5 *h* and *Hv* within species

To test whether *Hv* responds to height within species, we analysed individual records with both *h* and *Hv*, after removing duplicated *h*-*Hv* pairs and retaining species with ≥ 3 records (130 records, 22 species). We fitted a linear mixed model of log *Hv* on species-centred log *h*, with species as a random intercept, so that the slope reflects within-species variation only. *Hv* increased with *h* within species (slope = 0.34, 95% CI [0.17, 0.50]).

## REFERENCES

1. Ackerly, D. (2009). Conservatism and diversification of plant functional traits: Evolutionary rates versus phylogenetic signal. Proc. Natl. Acad. Sci. U.S.A., 106, 19699–19706.

2. Becker, P., Meinzer, F.C. & Wullschleger, S.D. (2000). Hydraulic limitation of tree height: a critique. Functional Ecology, 14, 4–11.

3. Binks, O., Cernusak, L.A., Liddell, M., Bradford, M., Coughlin, I., Carle, H., et al. (2022). Forest system hydraulic conductance: partitioning tree and soil components. New Phytologist, 233, 1667–1681.

4. Bollen, K.A. (1987). Total, Direct, and Indirect Effects in Structural Equation Models. Sociological Methodology, 17, 37.

5. Brodribb, T.J. & Jordan, G.J. (2008). Internal coordination between hydraulics and stomatal control in leaves. Plant Cell & Environment, 31, 1557–1564.

6. Brodribb, T.J. & McAdam, S.A.M. (2017). Evolution of the Stomatal Regulation of Plant Water Content. Plant Physiol., 174, 639–649.

7. Carminati, A., Javaux, M., Wankmüller, F.J.P. & Brodribb, T.J. (2026). Soils drive convergence in the regulation of vascular tension in land plants. Science, 391, 476–479.

8. Carpenter, B., Gelman, A., Hoffman, M.D., Lee, D., Goodrich, B., Betancourt, M., et al. (2017). *Stan* : A Probabilistic Programming Language. J. Stat. Soft., 76.

9. Choat, B., Cobb, A.R. & Jansen, S. (2008). Structure and function of bordered pits: new discoveries and impacts on whole-plant hydraulic function. New Phytologist, 177, 608–626.

10. Couvreur, V., Ledder, G., Manzoni, S., Way, D.A., Muller, E.B. & Russo, S.E. (2018). Water transport through tall trees: A vertically explicit, analytical model of xylem hydraulic conductance in stems. Plant Cell & Environment, 41, 1821–1839.

11. Fick, S.E. & Hijmans, R.J. (2017). WorldClim 2: new 1-km spatial resolution climate surfaces for global land areas. Intl Journal of Climatology, 37, 4302–4315.

12. Gabry, J., Češnovar, R., Johnson, A. & Bronder, S. (2025). cmdstanr: R Interface to “CmdStan.” Global Biodiversity Information Facility (GBIF). (2026). 10.15468/dl.sbpxnb.

13. Gonzalez-Voyer, A. & Von Hardenberg, A. (2014). An Introduction to Phylogenetic Path Analysis. In: Modern Phylogenetic Comparative Methods and Their Application in Evolutionary Biology (ed. Garamszegi, L.Z.). Springer Berlin Heidelberg, Berlin, Heidelberg, pp. 201–229.

14. Hadfield, J.D. & Nakagawa, S. (2010). General quantitative genetic methods for comparative biology: phylogenies, taxonomies and multi-trait models for continuous and categorical characters. Journal of Evolutionary Biology, 23, 494–508.

15. Halliwell, B., Holland, B.R. & Yates, L.A. (2025). Multi-response phylogenetic mixed models: concepts and application. Biological Reviews, 100, 1294–1316.

16. Hardenberg, A.V. & Gonzalez-Voyer, A. (2013). Disentangling evolutionary cause-effect relationships with phylogenetic confirmatory path analysis. Evolution, 67, 378–387.

17. Huber, B. (1928). Weitere quantitative Untersuchungen über das Wasserleitungssystem der Pflanzen. Jahrb Wiss Bot., 67, 877.

18. Jin, Y. & Qian, H. (2022). V.PhyloMaker2: An updated and enlarged R package that can generate very large phylogenies for vascular plants. Plant Diversity, 44, 335–339.

19. Kattge, J., Bönisch, G., Díaz, S., Lavorel, S., Prentice, I.C., Leadley, P., et al. (2020). TRY plant trait database – enhanced coverage and open access. Global Change Biology, 26, 119–188.

20. Lechthaler, S., Kiorapostolou, N., Pitacco, A., Anfodillo, T. & Petit, G. (2020). The total path length hydraulic resistance according to known anatomical patterns: What is the shape of the root-to-leaf tension gradient along the plant longitudinal axis? Journal of Theoretical Biology, 502, 110369.

21. Lin, Y.-S., Medlyn, B.E., Duursma, R.A., Prentice, I.C., Wang, H., Baig, S., et al. (2015). Optimal stomatal behaviour around the world. Nature Clim Change, 5, 459–464.

22. Liu, H., Gleason, S.M., Hao, G., Hua, L., He, P., Goldstein, G., et al. (2019). Hydraulic traits are coordinated with maximum plant height at the global scale. Sci. Adv., 5, eaav1332.

23. Liu, H., Xu, Q., He, P., Santiago, L.S., Yang, K. & Ye, Q. (2015). Strong phylogenetic signals and phylogenetic niche conservatism in ecophysiological traits across divergent lineages of Magnoliaceae. Sci Rep, 5, 12246.

24. Losos, J.B. (2008). Phylogenetic niche conservatism, phylogenetic signal and the relationship between phylogenetic relatedness and ecological similarity among species. Ecology Letters, 11, 995–1003.

25. Maherali, H., Pockman, W.T. & Jackson, R.B. (2004). Adaptive variation in vulnerability of woody plants to xylem cavitation. Ecology, 85, 2184–2199.

26. Martínez-Vilalta, J., Korakaki, E., Vanderklein, D. & Mencuccini, M. (2007). Below-ground hydraulic conductance is a function of environmental conditions and tree size in Scots pine. Functional Ecology, 21, 1072–1083.

27. Martínez-Vilalta, J., Prat, E., Oliveras, I. & Piñol, J. (2002). Xylem hydraulic properties of roots and stems of nine Mediterranean woody species. Oecologia, 133, 19–29.

28. Martin-StPaul, N., Delzon, S. & Cochard, H. (2017). Plant resistance to drought depends on timely stomatal closure. Ecol Lett, 20, 1437–1447.

29. McDowell, N., Phillips, N., Lunch, C., Bond, B. & Ryan, M. (2002). An investigation of hydraulic limitation and compensation in large, old Douglas-fir trees. Tree Physiology, 22, 763–774.

30. McDowell, N.G. & Allen, C.D. (2015). Darcy’s law predicts widespread forest mortality under climate warming. Nature Clim Change, 5, 669–672.

31. Meinzer, F.C. (2002). Co-ordination of vapour and liquid phase water transport properties in plants. Plant Cell Environ, 25, 265–274.

32. Meinzer, F.C., McCulloh, K.A., Lachenbruch, B., Woodruff, D.R. & Johnson, D.M. (2010). The blind men and the elephant: the impact of context and scale in evaluating conflicts between plant hydraulic safety and efficiency. Oecologia, 164, 287–296.

33. Mencuccini, M. (2003). The ecological significance of long-distance water transport: short-term regulation, long-term acclimation and the hydraulic costs of stature across plant life forms. Plant Cell & Environment, 26, 163–182.

34. Mencuccini, M. & Grace, J. (1996). Developmental patterns of above-ground hydraulic conductance in a Scots pine (Pinus sylvestris L) age sequence. Plant Cell & Environment, 19, 939–948.

35. Mencuccini, M. & Magnani, F. (2000). Comment on “Hydraulic limitation of tree height: a critique” by Becker, Meinzer & Wullschleger. Functional Ecology, 14, 135–137.

36. Mencuccini, M., Rosas, T., Rowland, L., Choat, B., Cornelissen, H., Jansen, S., et al. (2019). Leaf economics and plant hydraulics drive leaf : wood area ratios. New Phytol, 224, 1544–1556.

37. Murray, M., Soh, W.K., Yiotis, C., Batke, S., Parnell, A.C., Spicer, R.A., et al. (2019). Convergence in Maximum Stomatal Conductance of C3 Woody Angiosperms in Natural Ecosystems Across Bioclimatic Zones. Front. Plant Sci., 10, 558.

38. Muzzamal, M., Huang, J., Nielson, R., Sefton, M. & Triantafilis, J. (2018). Mapping Soil Particle-Size Fractions Using Additive Log-Ratio (ALR) and Isometric Log-Ratio (ILR) Transformations and Proximally Sensed Ancillary Data. Clays and clay miner., 66, 9–27.

39. Olson, M.E., Anfodillo, T., Rosell, J.A., Petit, G., Crivellaro, A., Isnard, S., et al. (2014). Universal hydraulics of the flowering plants: vessel diameter scales with stem length across angiosperm lineages, habits and climates. Ecology Letters, 17, 988–997.

40. Olson, M.E., Soriano, D., Rosell, J.A., Anfodillo, T., Donoghue, M.J., Edwards, E.J., et al. (2018). Plant height and hydraulic vulnerability to drought and cold. Proc Natl Acad Sci USA, 115, 7551–7556.

41. Pagel, M. (1999). Inferring the historical patterns of biological evolution. Nature, 401, 877–884.

42. Petit, G. & Anfodillo, T. (2009). Plant physiology in theory and practice: An analysis of the WBE model for vascular plants. Journal of Theoretical Biology, 259, 1–4.

43. Phillips, N., Bond, B.J., McDowell, N.G. & Ryan, M.G. (2002). Canopy and hydraulic conductance in young, mature and old Douglas-fir trees. Tree Physiology, 22, 205–211.

44. Poggio, L., De Sousa, L.M., Batjes, N.H., Heuvelink, G.B.M., Kempen, B., Ribeiro, E., et al. (2021). SoilGrids 2.0: producing soil information for the globe with quantified spatial uncertainty. SOIL, 7, 217–240.

45. R Core Team. (2024). R: a language and environment for statistical computing (manual). R Foundation for Statistical Computing, Vienna, Austria.

46. Revell, L.J. (2024). phytools 2.0: an updated R ecosystem for phylogenetic comparative methods (and other things). PeerJ, 12, e16505.

47. Ryan, M.G., Phillips, N. & Bond, B.J. (2006). The hydraulic limitation hypothesis revisited. Plant Cell & Environment, 29, 367–381.

48. Ryan, M.G. & Yoder, B.J. (1997). Hydraulic Limits to Tree Height and Tree Growth. BioScience, 47, 235–242.

49. Sabot, M.E.B., De Kauwe, M.G., Pitman, A.J., Medlyn, B.E., Verhoef, A., Ukkola, A.M., et al. (2020). Plant profit maximization improves predictions of European forest responses to drought. New Phytol, 226, 1638–1655.

50. Sack, L. & Holbrook, N.M. (2006). Leaf hydraulics. Annu. Rev. Plant Biol., 57, 361–381.

51. Sanchez-Martinez, P., Ackerly, D.D., Martínez-Vilalta, J., Mencuccini, M., Dexter, K.G. & Dawson, T.E. (2024). A framework to study and predict functional trait syndromes using phylogenetic and environmental data. Methods Ecol Evol, 15, 666–681.

52. Sanchez-Martinez, P., Martínez-Vilalta, J., Dexter, K.G., Segovia, R.A. & Mencuccini, M. (2020). Adaptation and coordinated evolution of plant hydraulic traits. Ecol Lett, ele.13584.

53. Scheffer, M., Xu, C., Hantson, S., Holmgren, M., Los, S.O. & Van Nes, E.H. (2018). A global climate niche for giant trees. Global Change Biology, 24, 2875–2883.

54. Shipley, B. (2009). Confirmatory path analysis in a generalized multilevel context. Ecology, 90, 363–368.

55. Smith, S.A. & Brown, J.W. (2018). Constructing a broadly inclusive seed plant phylogeny. American J of Botany, 105, 302–314.

56. Tao, S., Guo, Q., Li, C., Wang, Z. & Fang, J. (2016). Global patterns and determinants of forest canopy height. Ecology, 97, 3265–3270.

57. Thorson, J.T. & Van Der Bijl, W. (2023). phylosem: A fast and simple R package for phylogenetic inference and trait imputation using phylogenetic structural equation models. Journal of Evolutionary Biology, 36, 1357–1364.

58. Turek, M.E., Poggio, L., Batjes, N.H., Armindo, R.A., De Jong Van Lier, Q., De Sousa, L., et al. (2023). Global mapping of volumetric water retention at 100, 330 and 15 000 cm suction using the WoSIS database. International Soil and Water Conservation Research, 11, 225–239.

59. Van Der Bijl, W. (2018). *phylopath* : Easy phylogenetic path analysis in R. PeerJ, 6, e4718.

60. Vehtari, A., Gelman, A., Simpson, D., Carpenter, B. & Bürkner, P.-C. (2021). Rank-Normalization, Folding, and Localization: An Improved R^ for Assessing Convergence of MCMC (with Discussion). Bayesian Anal., 16.

61. Westoby, M., Yates, L., Holland, B. & Halliwell, B. (2023). Phylogenetically conservative trait correlation: Quantification and interpretation. Journal of Ecology, 111, 2105–2117.

62. Whittaker, R. (1975). Communities and ecosystems. 2. ed., 1. print. Macmillan [u.a.], New York.

63. Xu, H., Wang, H., Prentice, I.C., Harrison, S.P., Rowland, L., Mencuccini, M., et al. (2026). Global variation in the ratio of sapwood to leaf area explained by optimality principles. New Phytologist, 250, 181–193.

64. Zaehle, S. (2005). Effect Of Height On Tree Hydraulic Conductance Incompletely Compensated By Xylem Tapering. Functional Ecology, 19, 359–364.

65. Zhao, Y., Mencuccini, M., Acuña-Míguez, B., Adet, L., Amin, S., Anadon-Rosell, A., et al. (2026). Global dataset of plant water transport capacity. Sci Data.

66. Zizka, A., Silvestro, D., Andermann, T., Azevedo, J., Duarte Ritter, C., Edler, D., et al. (2019). COORDINATECLEANER : Standardized cleaning of occurrence records from biological collection databases. Methods Ecol Evol, 10, 744–751.

67. Zomer, R.J., Xu, J. & Trabucco, A. (2022). Version 3 of the Global Aridity Index and Potential Evapotranspiration Database. Sci Data, 9, 409.

